# Genotype-to-phenotype mapping of somatic clonal mosaicism via single-cell co-capture of DNA mutations and mRNA transcripts

**DOI:** 10.1101/2024.05.22.595241

**Authors:** Dennis J Yuan, John Zinno, Theo Botella, Dalia Dhingra, Shu Wang, Allegra Hawkins, Ariel Swett, Jesus Sotelo, Ramya Raviram, Clayton Hughes, Catherine Potenski, Akira Yokoyama, Nobuyuki Kakiuchi, Seishi Ogawa, Dan A Landau

**Affiliations:** New York Genome Center, New York, NY; Division of Hematology and Medical Oncology, Department of Medicine, Weill Cornell Medicine, New York, NY; Sandra and Edward Meyer Cancer Center, Weill Cornell Medicine, New York, NY; Mission Bio, San Francisco, CA; Institute for the Advanced Study of Human Biology, Kyoto University, Kyoto, Japan; Department of Medical Oncology, Kyoto University, Kyoto, Japan; The Hakubi Center for Advanced Research, Kyoto University, Kyoto, Japan

## Abstract

Somatic mosaicism is a hallmark of malignancy that is also pervasively observed in human physiological aging, with clonal expansions of cells harboring mutations in recurrently mutated driver genes. Bulk sequencing of tissue microdissection captures mutation frequencies, but cannot distinguish which mutations co-occur in the same clones to reconstruct clonal architectures, nor phenotypically profile clonal populations to delineate how driver mutations impact cellular behavior. To address these challenges, we developed <u>s</u>ingle-<u>c</u>ell <u>G</u>enotype-to-<u>P</u>henotype sequencing (scG2P) for high-throughput, highly-multiplexed, single-cell joint capture of recurrently mutated genomic regions and mRNA phenotypic markers in cells or nuclei isolated from solid tissues. We applied scG2P to aged esophagus samples from five individuals with high alcohol and tobacco exposure and observed a clonal landscape dominated by a large number of clones with a single driver event, but only rare clones with two driver mutations. *NOTCH1* mutants dominate the clonal landscape and are linked to stunted epithelial differentiation, while *TP53* mutants and double-driver mutants promote clonal expansion through both differentiation biases and increased cell cycling. Thus, joint single-cell highly multiplexed capture of somatic mutations and mRNA transcripts enables high resolution reconstruction of clonal architecture and associated phenotypes in solid tissue somatic mosaicism.

## Introduction

Somatic evolution involves ongoing genetic diversification in human tissues. In the accelerated evolutionary process of cancer, clonal diversification allows malignancies to enhance their fitness and overcome therapeutic challenges^1–3^. Clonal mosaicism (CM) has more recently been demonstrated across normal^4–6^ and diseased non-malignant^7–11^ human tissues. In a striking example, genomic profiling of phenotypically normal esophagus (PNE) has uncovered the existence of a diverse landscape of driver-mutated clones^4,12^. PNE tissue is colonized by clones harboring mutations in cancer-associated driver genes, with clonal expansion increasing with age. While driver-mutated clones account for less than 10% of the epithelium for individuals in their 20s, such clones can replace up to 80% of the epithelium in individuals older than 60^4,12,13^. Despite the encroachment of a large number of diverse clones, PNE tissue maintains histological integrity, cell type composition, and function^14,15^. Driver mutations in PNE overlap with those in esophageal squamous cell carcinoma (ESCC), most notably *NOTCH1, TP53, NOTCH2, NOTCH3*, and *FAT1*^4,12^. However, some driver mutations appear at a higher frequency in normal tissue compared to ESCC, indicating that some clones can be negatively selected against in carcinogenesis. In the aging esophagus, *NOTCH1* mutants have been shown to outcompete tumor cells, but maintain tissue integrity similar to wild-type cells^14,15^. The ubiquity of somatic clonal expansions with aging and disease mark somatic mosaicism as a key frontier in human genetics.

The study of non-malignant CM with the prevailing strategy of microdissection followed by bulk sequencing has left two major gaps in our understanding of somatic mosaicism. First, bulk sequencing has limited ability to resolve clonal hierarchies of observed mutations, which broadly align with either a nested structure (two driver mutations within the same clone) or a branched structure (driver mutations impact sister clones)^13^. It is therefore largely unknown whether non-malignant tissue clones contain multiple driver mutations, or if instead driver mutations affect distinct clones, a gap that requires highly multiplexed mutational capture at single-cell resolution. Second, to date studies have focused exclusively on DNA sequencing of somatic mutations, without capturing their phenotypic impact in primary human samples. To explore the phenotypic effects of somatic mutations, we require single-cell multi-modality technologies that can link genotype and phenotypic readouts (e.g., transcriptome or chromatin accessibility) at the single-cell resolution.

In blood mosaicism, the ability to link genotype to phenotype in single cells revealed that CM drivers often disrupt differentiation hierarchies. These studies rely on either plate-based methods^16–18^ where throughput is limited, especially considering the small clone size in CM, or droplet-based single-cell assays (such as Genotyping of Transcriptomes^19,20^) that enable higher throughput genotyping together with transcriptome profiling. Notably, these investigations have exclusively focused on clonal outgrowths within the hematopoietic system, as ease of blood draw allows for prior genotyping of bulk samples to direct limited targeted mutational capture with single-cell multimodal sequencing methods. Viably frozen whole cells from blood additionally ensures higher mRNA content in contrast to single-nucleus approaches that are often required for the study of archival solid tissues, facilitating mutational capture of transcribed genes in scRNAseq. In contrast, genotype-to-phenotype mapping in solid tissue CM is more challenging. For example, in PNE, mutations often distribute through large genomic territories, hindering isolated hotspot capture genotyping approaches. Moreover, clones are comparatively small and spatially segregated, limiting the ability to have prior knowledge about the specific mutation before the application of single-cell genomics. Archival solid tissue samples require nuclei extraction, resulting in lower mRNA content. As such, application of single-cell genotype-to-phenotype mapping of CM in solid tissue has been limited, resulting in a large knowledge gap regarding the phenotypic changes in somatic mosaicism of non-hematopoietic tissues.

To address this challenge, we developed <u>s</u>ingle-<u>c</u>ell <u>G</u>enotype-to-<u>P</u>henotype sequencing (scG2P), a single-cell approach for the highly multiplexed capture of multiple recurrently mutated regions in driver genes to decipher mosaicism in solid tissue, while elucidating cell states with an mRNA readout. The high-throughput microfluidic methodology enables the profiling of thousands of cells required for the capture of smaller clones in solid tissue mosaicism. Through cell mixing analysis, we demonstrate accurate co-capture of 118 genomic amplicons and 56 mRNA transcripts that serve as phenotypic markers. We applied targeted DNA and mRNA co-capture and profiled over 5,000 single nuclei isolated from aged PNE samples from five donors with high tobacco and alcohol exposure, enabling detection of somatic variants across six driver genes (*NOTCH1, TP53, NOTCH2, NOTCH3, FAT1, PPM1D*). We resolved the clonal architecture of PNE, with the majority of clones driven by single driver mutations, and rare clones with two driver mutations. Using the matched transcriptional information, we assigned cells with epithelial differentiation stages and cell cycle scores to define varying driver-specific phenotypes. For example, while frequent *NOTCH1*-mutated clones show decreased differentiation as the leading fitness-enhancing mechanism, *TP53* mutant clones have enhanced fitness through both decreased differentiation and increased proliferation. Collectively, these results provide key insights into somatic evolution in normal solid tissue, uncovering the clonal architecture at high resolution, as well as a matrix of genotype-to-phenotype associations, delivering a novel framework for deciphering the functional consequences of somatic mutations in solid tissues.

## Results

### Co-capture of highly multiplexed somatic mutation profiling and targeted mRNAs in single cells

Capturing the diversity of somatic mutants in solid tissue requires targeting a range of loci along driver genes. We utilized a single-cell DNA sequencing (scDNAseq) technology developed to capture mutational hotspots that has been used to assess clonal structure in AML^21,22^, chronic lymphocytic leukemia^23^ (CLL), and breast cancer^24^. This microfluidic platform uses a double encapsulation strategy, where the first encapsulation releases DNA content through cell lysis, followed by targeted amplification, and a second encapsulation step that adds cell barcodes to targeted amplicons. To further link genotypes to cell states, we added capture of mRNA transcripts of interest together with DNA amplicons in the same cell. To develop scG2P (single-cell Genotype to Phenotype), we modified the assay for reverse transcription of mRNA targets during the first encapsulation step, adding capture handles to cDNA amplicons for downstream barcoding. During the second encapsulation step, we add cell barcodes to the DNA and RNA amplicons, followed by multiplexed PCR amplification. To differentiate amplified transcripts from off-target capture of gDNA, we designed RNA capture primers to cover exon-exon junctions (**Fig. 1A**). Barcoded RNA and DNA amplicons are separated with streptavidin bead pulldown post emulsion amplification for separate library preparation. The final barcoded DNA and RNA amplicon libraries allow us to link somatic mutations with RNA signals through shared cell barcodes.

**Figure 1.**
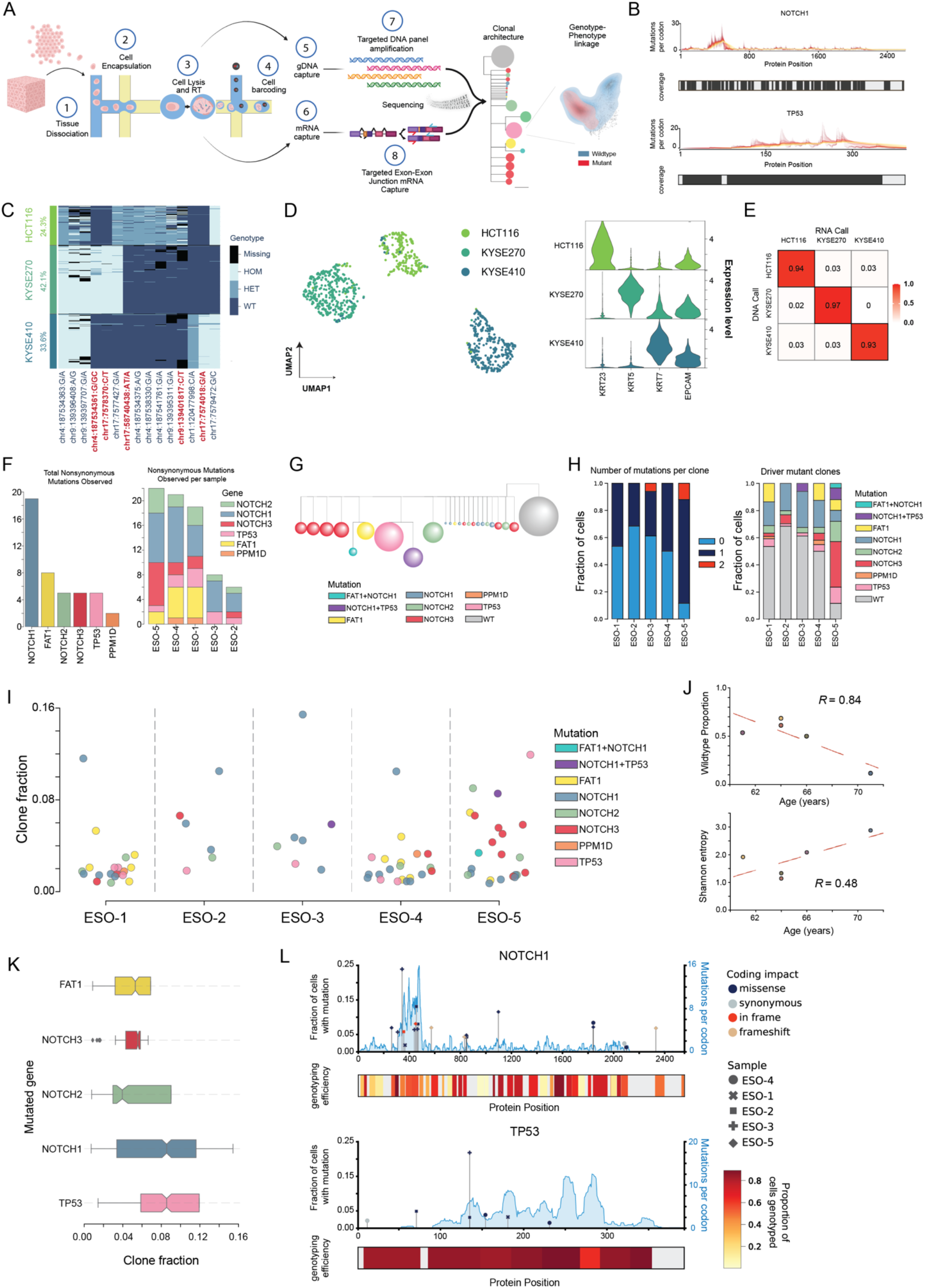
Targeted capture of mutation hotspots and RNA in single cells. **A**. Schematic representation of the scG2P workflow. Dissociated cells or nuclei undergo cell lysis and targeted reverse transcription (RT), followed by barcode addition and targeted loci amplification in two sequential encapsulations. DNA and RNA amplicons are separated by streptavidin bead capture for separate library preparation. This workflow combines DNA mutation hotspot capture to reconstruct clonal architecture with exon-exon capture of RNA targets for single-cell genotype to phenotype linkage. **B**. Tiling amplicon design and mutation frequency across *NOTCH1* and *TP53* genes. The number of mutations per codon, as previously reported from bulk sequencing of the esophagus (Yokoyama et al.), is displayed across the protein positions. The color scale represents the size of the rolling window used for calculating mutations per codon, with yellow indicating a 150 bp window and red indicating a 4 bp window. Black bars represent the presence of an amplicon covering the loci. Amplicon designs for *NOTCH2, NOTCH3*, and *FAT1* are provided in Supplementary Figure 1. **C**. Heatmap of filtered variants detected in mixing study. Cell lines HCT116, KYSE270, and KYSE410 were mixed and processed using scG2P. The heatmap displays the detected filtered variants for each individual cell, clustered by cell line based on the variant allele frequencies (VAFs) of the DNA variants. Variants annotated in red were independently validated using whole-exome sequencing data from the Cancer Cell Line Encyclopedia (CCLE) database. The genotype of each variant is indicated as homozygous (HOM), heterozygous (HET), wild-type (WT), or missing. **D**. (Left) RNA expression-based uniform manifold approximation projection (UMAP) for cell line mixing experiment. HCT116 (n = 234 cells), KYSE270 (n = 403 cells), and KYSE410 (n = 355 cells) are colored according to their assigned cell line identity, determined by k-means clustering of the variant allele frequencies of DNA variants. (Right) Violin plots displaying the RNA expression levels (center log ratio) of four marker genes (*KRT23, KRT5, KRT7*, and *EPCAM*) across the three cell lines. **E**. Confusion matrix comparing RNA-based clustering labels (predicted) and DNA-based clustering labels (ground truth) of the cell line mixing study. The matrix displays the percentage of cells assigned to each cell line based on RNA expression profiles compared to the ground truth DNA-based assignments. Diagonal elements represent correctly classified cells, while off-diagonal elements indicate misclassifications. The mean accuracy across all cell lines is 0.95. **F**. Nonsynonymous mutations captured across driver genes, stratified by gene and sample. (Left) Bar plot displaying the number of nonsynonymous mutations detected in each driver gene across all samples. (Right) Bar plot showing the contribution of nonsynonymous mutations by each driver gene for each patient sample. **G**. Clonal structure of ESO-5. Terminal nodes represent distinct subclones, with node sizes proportional to the relative mutant cell fraction of each subclone. Branch lengths are scaled to reflect the acquisition of a single mutation. **H**. Mutational burden per clone and distribution of clonal mutations across driver genes. (Left) Fraction of cells harboring 0, 1, or 2 mutations per clone for cells passing the 50% genotyping completeness threshold. (Right) Fraction of cells with mutation in indicated driver gene, or combination of mutations in driver genes. **I**. Clones detected across all donors. Circles represent clones called in each donor, colored by driver mutations detected in the clone, and y-axis representing the fraction of total cells each clone makes up in the sample. **J**. (Top) Scatter plot illustrating the Pearson correlation between the proportion of wild-type cells and donor age (R = 0.84). (Bottom) Scatter plot illustrating the Pearson correlation between clonal diversity, computed by the Shannon entropy index, and donor age (R = 0.48). Each point represents a donor, and the lines represent the linear fit. **K**. Clone fractions of cells with single driver mutation. Box plot display the distribution of mean mutant cell fractions of clones for each mutated gene across all donors (Center line, median; box, IQR; whisker, 1.5*IQR). Only cells with single mutations were included. **L**. Fraction of cells with mutation in *NOTCH1* and *TP53* genes. The cell fractions (left y-axis) of detected variants for each sample are plotted along the protein positions of *NOTCH1* and *TP53*. Each variant is annotated with its coding impact (missense, synonymous, in frame, or frameshift mutation). The mutant cell fraction plot is overlaid with the hotspot mutation density, as shown in **B**, representing the number of mutations per codon (right y-axis) as previously reported by Yokoyama et al. from bulk sequencing of the esophageal microdissections. The genotyping efficiency is displayed below each plot using a color scale that indicates the proportion of cells genotyped at each locus. Note that fraction of cells with mutation is represented by the clone fraction in all single-mutant clones, whereas in 2 out of 5 donors with detected double mutant clones, the mutant fraction is the sum of the parent clone and double mutant subclone fractions.

To test the ability of this technology to study human somatic mosaicism, we focused on PNE, where somatic mosaicism was shown to be pervasive in aging donors. We designed amplicon panels that capture frequently mutated sites across *NOTCH1, NOTCH2, NOTCH3, PPM1D, FAT1*, and *TP53*, previously identified as driver genes with the highest mutational burdens observed in PNE^4,12^ (**Supplementary Table 1**). Notably, while previous scDNAseq panels have largely focused on isolated mutational hotspots, we designed amplicons tiling across entire gene bodies to better characterize the mutational landscape. *NOTCH1, NOTCH2, NOTCH3, TP53* have known key roles in keratinocyte differentiation in both esophageal epithelium and skin, and contain positively selected mutations in both aging esophagus and sun-exposed skin^25–27^. *NOTCH1* mutations have been well characterized in conferring competitive advantage to mutant cells over wild-type cells without disrupting the epithelium structure^14^. Although G-C content or primer interactions resulted in variable capture efficiency of tiled regions, the amplicons covered regions that encompassed over 400 previously reported mutations in PNE in >21Kb of genomic DNA across the six driver genes (**Supplementary Fig. 1A**), allowing us to genotype cells across these recurrently mutated regions (**Fig. 1B, Supplementary Fig. 1B**).

To define informative transcripts for the design of an mRNA panel to assign cell states, we performed single-cell RNA sequencing and spatial transcriptomics on PNE biopsy samples. Human esophageal tissue is composed of a layer of basal epithelial cells farthest from the lumen that serves as a reservoir of progenitor cells. The next layers are composed of proliferating suprabasal cells that actively migrate towards the lumen, with an increasingly differentiated phenotype marked by becoming flattened and forming a tight barrier across the lumen. Differentiated cells shed as cells migrate, maintaining homeostasis and cell density^14,28^. We procured endoscopy punch biopsy samples from older individuals with high alcohol and/or tobacco exposure (**Supplementary Information**). We performed scRNAseq on dissociated single cells from three donors with 10x Genomics Chromium assays and annotated cell types based on cluster similarity and marker genes (**Supplementary Fig. 2A, 2B, 2C**) from an esophageal single-cell reference dataset^29,30^. These data showed that our tissue sampling methodology in aged individuals captured the established trajectory of epithelial cell differentiation, spanning from the early basal state through the differentiating suprabasal state, to the differentiated cells in the upper and stratified regions (**Supplementary Fig. 2B, 2C**). To determine the spatial segregation of populations within the biopsied area, we also performed spatial transcriptomics (ST) utilizing the 10x Visium platform to assay sections from optimal cutting temperature compound (OCT)-embedded biopsy punches from three donors (**Supplementary Information**). Using annotations from the single-cell datasets, we categorized spatial barcodes as clusters of cell types and observed regions composed of basal, suprabasal, and differentiated epithelial cell types (**Supplementary Fig. 2D**). Trajectory analysis of ST barcodes recapitulated these differentiation transcriptional signatures, comprising markers of basal cells (*KRT5, KRT15*), suprabasal cells (*KRT13, KRT4*), and differentiated cells (*S100A9, SPRR3*) (**Supplementary Fig. 2E, 2F**). We additionally confirmed *TP63, SOX2*, and *COL17A1* as regulators of epithelial differentiation in basal and suprabasal cells (**Supplementary Fig. 2F**), as previously reported^29,30^. Notably, the scRNAseq and ST data revealed similar esophageal tissue structure across aged donors with PNE, underscoring the need for multi-modal genotype single-cell information to determine mutant and wild-type cell composition and phenotype at high resolution. Using these single-cell and spatial datasets, we designed a targeted mRNA panel of 56 markers (**Supplementary Table 2**; **Methods**) to enable identification of the cell types, stages of epithelial differentiation, and markers of stemness and proliferation in mutant vs. wild-type cells. Using this set of mRNA targets alone recapitulates the cell types observed from scRNAseq datasets (**Supplementary Fig. 2G**).

To validate the performance of scG2P, we performed cell line mixing with two esophageal cell lines (KYSE-270 and KYSE-410) and a colon cell line (HCT-116). Valid cell barcodes were determined by read depth and coverage uniformity across the DNA panel (Methods), and variants were filtered based on read depth, allelic frequency, and genotyping quality (**Supplementary Table 3**), as previously described^21–24^. Cell lines were identified by clustering based on variant allele frequencies (VAFs) of filtered variants (**Fig. 1C**). We identified nine single-nucleotide variants (SNVs) that were private to individual cell lines. Five of these were variants previously identified by whole-exome sequencing (WES) in the Cancer Cell Line Encyclopedia (CCLE) database (**Fig. 1C**). The remaining four mutations (one intronic, three coding) likely represent mutations acquired by individual cell lines during culture (**Supplementary Fig. 3A**). We also detected in each cell line three additional shared germline variants that have high population frequency according to the GnomAD database (**Supplementary Fig. 3A, Supplementary Table 3**). Genotyping accuracy, calculated as the percentage of the correct WES variant call for each cell according to cell line assignment, was 92% (SEM: 3.64%) (**Supplementary Fig. 3B**). We compared VAFs of the known WES and phased variants (Methods) to determine distributions of false positives and allelic dropout (ADO) rates (**Supplementary Fig. 3C**), and estimated a median ADO of <10% (**Supplementary Fig. 3D**). Comparing amplicon performance, we observed that amplicon G-C content decreased amplicon capture efficiency, most notably impacting some of the G-C rich *NOTCH1* amplicons (**Supplementary Fig. 3E**). Additionally, shorter insert length decreased amplicon capture efficiency, but this effect was more dispersed across driver genes (**Supplementary Fig. 3F**). We also estimated known copy number alterations (CNAs) of the cell lines, but only successfully detected two of the four known events, likely due to the low number of amplicons covered per chromosome region (**Supplementary Fig. 3G)**. Therefore, we did not pursue CNA analysis using this current targeted panel in our primary samples. Ultimately, our DNA panel performed similarly to previous scDNAseq panels^21–24^ in terms of genotyping efficiency and cells recovered, with the addition of mRNA capture (**Supplementary Table 4**).

To integrate mRNA data, we aligned mRNA amplicons and removed reads that did not align to primer probe positions. To distinguish between genomic DNA contamination and captured transcript, we filtered out reads with alignment to intronic regions (**Supplementary Fig. 4A**). We generated a cell by gene counts matrix using the filtered reads for downstream analysis and observed three clusters representing each of the expected cell lines (**Fig. 1D, Supplementary Table 5**). We note that RNA reads that passed filtering were strongly correlated with DNA reads in the cell line mixing with whole cells, but were not as strongly correlated in isolated nuclei from tissue samples (**Supplementary Fig. 4B)**. We assigned cell line identity based on expression level of cell line-specific marker genes (*KRT5, KRT7, KRT23*) in each cluster (**Fig. 1D)**. To determine genotype-phenotype linkage accuracy, we used genotype assignments as ground truth compared to RNA-based clustering, achieving an accuracy of 0.95 (**Fig. 1E**). We then compared the pseudo-bulk mRNA expression levels in our data to bulk RNAseq expression levels from the CCLE dataset and obtained Spearman’s correlation of 0.72-0.76 across the cell lines (**Supplementary Fig. 4C)**. We then clustered the three cell lines based on panel RNA expression and annotated the UMAPs with three ground truth variants called via WES that distinguish the cell types, showing accurate cell type assignment (**Supplementary Fig. 4D**). Collectively, these data demonstrate that our approach enables highly multiplexed capture of mutations and targeted mRNA profiling in single cells, with high accuracy in assessing genotype to phenotype linkage.

### Human normal esophageal epithelium is largely composed of somatic clones with single driver mutations

Bulk micro-dissection targeted sequencing has revealed that the esophageal epithelium is a mosaic of somatically mutated cells; yet the precise clonal structure, including clone size and mutation complexity, cannot be fully resolved in bulk sequencing. To define the clonal architecture of somatic mosaicism of PNE, as well as link somatic mutations with cellular phenotypes, we isolated single nuclei from OCT-embedded punch biopsies from five donors **(Supplementary Information)** as input for scG2P (2 mm^2^ biopsy punches, biopsies approximately 10 mm apart, 2 biopsies used per donor). Barcoded amplicons were used to call cells and variants, followed by genotyping and clonal reconstruction (**Supplementary Fig. 5A)**. We recovered and used as input between 40,000-120,000 (median: 50,000) nuclei per pair of biopsy punches and called between 713-1,747 (median: 1,105) cells per sample (Methods; **Supplementary Fig. 5B**). Mean sequencing depth of amplicons across the six driver genes ranged from 87X-270X per cell (**Supplementary Fig. 5C, 5D**), with 80% or more of amplicons in each driver gene achieving sufficient depth for genotyping (**Supplementary Fig. 5E**). After variant calling, we filtered for variants based on minimum read depth and genotype quality, and detected between 6-22 (median: 19) nonsynonymous somatic variants per sample (**Fig. 1F, Supplementary Table 6**), with *NOTCH1* mutations representing the majority of mutations across all samples (**Fig 1F**), as expected from previous bulk profiling of PNE^4,12^. Along with mutations in *NOTCH1*, mutations in *TP53 and NOTCH2*, previously identified as the second and fifth most frequently mutated genes, respectively, in the PNE from bulk profiling^12^, were detected in all samples (**Fig. 1F**).

To account for allelic dropout, we filtered cells based on their genotyping completeness, which is the percentage of variant loci detected in a cell of all variant loci detected in the donor. We set the minimum genotyping completeness at 50%, so that every cell has a call for at least 50% of the variants in that sample (Methods). While increased genotyping completeness stringency lowered the number of cells with high dropout, this also decreased cell numbers for analysis power (**Supplementary Fig. 6A**). We used VAFs as features to assign cells to genotype clusters based on density-based clustering, then merged similar clusters to obtain clonal assignment (Methods, **Supplementary Fig. 5A**). As comparison, we tested a second clonal inference approach by adapting a previously developed reinforcement learning model applied to similar scDNAseq data^21^. We note that these approaches were specifically developed for the use of single-cell genotyping panels for clonal reconstruction, and we used both methods to construct clonal structure, observing the same structure across all samples (**Fig. 1G, Supplementary 6B**). Of note, we observed that the median genotyping completeness of cells within a clone did not affect clone size estimates, as there was no correlation between the two measures (**Supplementary Fig. 6C**). At 50% genotyping completeness, we found that between 31-89% (median: 45%) of captured cells had at least one driver mutation across donors (**Fig. 1H**). The majority of mutant cells had only a single detected mutation and clones with two mutations were observed in only two samples (**Fig. 1I**). To determine whether mutant cell fractions could be underestimated due to dropout, we reanalyzed the dataset with a more stringent filtering of >80% genotyping completeness. At this higher threshold, the number of mutant cells increased slightly to 34-92% (median: 52%) (**Supplementary Fig. 6D**). However, the same double mutant clones and parent clones (harboring only one mutation out of the two found in the double mutant clones) in ESO-5 were identified at similar proportions compared to clonal analysis with 50% genotyping completeness (**Supplementary Fig. 6E**), suggesting that dropout does not drastically hinder our ability to detect double mutant clones.

Our single-cell data revealed that the majority of clones detected were driven by a single driver mutation. Thus, despite the high rates of driver mutations detected in PNE overall, these mutations are driving different clonal populations. To quantify the size and diversity of clonal populations, we quantified clone fraction (CF), which is the proportion of cells found within a clonal assignment out of all cells in that sample. The heterogeneous clonal landscape was reflected in clonal size through a high diversity in CF, where maximum CF ranged from 11-16% (median: 12%) across samples (**Fig. 1I**). *NOTCH1-*mutated clones represented the largest detected clone in four out of five donors, comprising between 10-16% of cells per sample (**Fig. 1I, Supplementary Fig. 6B**), with many additional *NOTCH1*-mutated clones (between 3-8 clones per sample) uncovered at smaller clone fractions. Although less frequent than *NOTCH1*-mutated clones, at least one *TP53*-mutated clone was detected in all samples (between 1-3 clones per sample), and was the largest clone in one sample (ESO-5). Interestingly, the oldest donor we sampled (ESO-5, 71 years old) had a large number of mutant cells harboring *NOTCH3* mutations (**Fig. 1I**), while *NOTCH3* mutations were completely absent in ESO-3, and were 3-to 5-fold less frequent than *NOTCH1* mutations in the remaining three donors. This variation in *NOTCH3* mutation rates has been previously reported^4^, and may reflect not only the effect of age, but differences in exposure to environmental factors or in genetic background as well. Notably, we found that the proportion of wild-type cells decreased with age (**Fig. 1J**, top) with a reciprocal increase in clonal diversity, as measured by Shannon Index that accounts for number of clones and their relative abundance per sample (**Fig. 1J**, bottom), suggesting that the age-associated increased mutation burden observed in the esophagus is due to the formation of large numbers of mutant clones of varying sizes, instead of a large dominant clone. To directly assess the level of growth advantage imparted by mutations across clones, we stratified clone sizes based on driver mutation and observed that *NOTCH1* and *TP53* mutations resulted in larger clone fractions than those with *NOTCH2, NOTCH3*, or *FAT1* mutations (**Fig. 1K)**. The mutations driving these larger expansions occurred in previously reported high mutation frequency regions^12^ (**Fig. 1L**), further suggesting a relationship between selection of clones with *NOTCH1* and *TP53* mutations and their larger sizes compared not only to wild-type cells but also to other driver mutants. Together, our observation of increased clonal diversity during aging (**Fig. 1J**) indicates that there is continued expansion of multiple clonal populations that compete in aging tissue. These results are consistent with the model where neighboring mutant clones of approximately similar fitness compete in the spatially ordered esophageal epithelium to maintain tissue homeostasis over time^14^, resulting in a lack of dominant clones that have a high enough selective advantage to drive down clonal diversity with age.

Bulk sequencing of PNE microdissections has previously revealed examples of branching evolution of subclones after an initial large clonal expansion from a driver mutation. This analysis in bulk is only applicable for clones that are sufficiently large to apply assumptions such as pigeonhole principle^4,31^. To investigate the hierarchical architecture of subclones, we took advantage of the resolution of our single-cell data to detect and quantify double-mutant clones that would be too small for the pigeonhole principle to apply, enabling us to identify double mutants in two samples. In ESO-3, we detected a clone with co-mutations in *NOTCH1* and *TP53* (CF = 6%) and its related *NOTCH1*-mutant parent clone (CF = 16%) (**Fig. 1I**). In ESO-5, we detected a clone with mutations in *TP53* and *NOTCH1* (CF = 8%) and its related *TP53* parent clone (CF = 12%) (**Fig. 1I**). In addition, we observed a second clone with co-mutations in *NOTCH1* and *FAT1* (CF = 4%) with the related *FAT1*-mutant clone (CF = 6%) (**Fig. 1I**). Notably, both the *NOTCH1*-mutated clone in ESO-3 and the *TP53*-mutated clone in ESO-5 that produced the double mutant subclones were the largest detected clones in those donors (**Fig. 1I**), suggesting ongoing evolution in larger clones. Altogether, we demonstrate that our scDNAseq strategy allows us to determine clonal structure in esophageal clonal mosaicism, particularly in resolving clones with double mutations and unveiling a highly mutated esophageal epithelium dominated by a large number of single mutant clones driven primarily by *NOTCH1* and *TP53* mutations.

### Genotype-phenotype mapping of somatic mutant clones in the PNE

Our scDNAseq strategy revealed a large number of clones, of diverse sizes, driven primarily by single driver mutation in the aging PNE. With the associated epithelial mRNA panel, we sought to further decompose cell type biases to gain insights into possible mechanisms supporting selection advantage in expanded clones. We annotated the cell clusters using canonical marker genes, identifying basal (*COL17A1+, TP63+, KRT15+*), basal DCN+ (*DCN+, COL17A1+, TP63+, KRT15+*), basal proliferative (*COL17A1+, TP63+, KRT15+, UBE2C+, MKI67+, TOP2A+*), suprabasal (*TP63+, COL17A1-*), suprabasal KLF4+ (*TP63+, COL17A1-, KLF4+)*, mature suprabasal (*SPRR3+, TP63-*), and fibroblast (*COL1A2+, DCN+*) cells (**Fig. 2A, Supplementary Table 7**). Within the basal cell population, we identified a previously reported quiescent population marked by *DCN* expression^30^ and a proliferative population marked by cell cycling gene expression (**Fig. 2A, 2B**). We computed a cycling score using our panel’s cycling genes (*UBE2C, MKI67, TOP2A*) and a differentiation score by using the difference between early epithelial score (*KRT15, TP63, COL17A1, KRT14*) and late epithelial module score (*SPRR3, S100A8, S100A9, KRT4, KRT13*) (**Fig. 2B, Supplementary Table 8**). These module scores provided a continuous scale that enabled us to place individual cells along a continuum of cell cycle and differentiation. In addition, we mapped epithelial cells along diffusion components (**Fig. 2C**) and projected the clonal proportions across driver mutation along pseudotime deciles reflecting differentiation state, enabling visualization of cell state changes per donor (**Fig. 2D, Supplementary Fig. 7**). We observed that clones with mutations in *TP53* (ESO-2, ESO-5), *NOTCH1*+*TP53* mutant (ESO-3), and *TP53*+*NOTCH1* and *FAT1*+*NOTCH*1 double mutants (ESO-5) were enriched in earlier differentiation states (**Fig. 2D, Supplementary Fig. 7**). Through calculating the differential contribution of the mutant clones to cell populations across the stages of epithelial differentiation, we can assess differentiation bias as a potential mechanism for clonal fitness advantage, providing a quantitative framework to define how driver mutations alter cell state proportions across PNE differentiation **(Fig. 2D, Supplementary Table 9)**.

**Figure 2.**
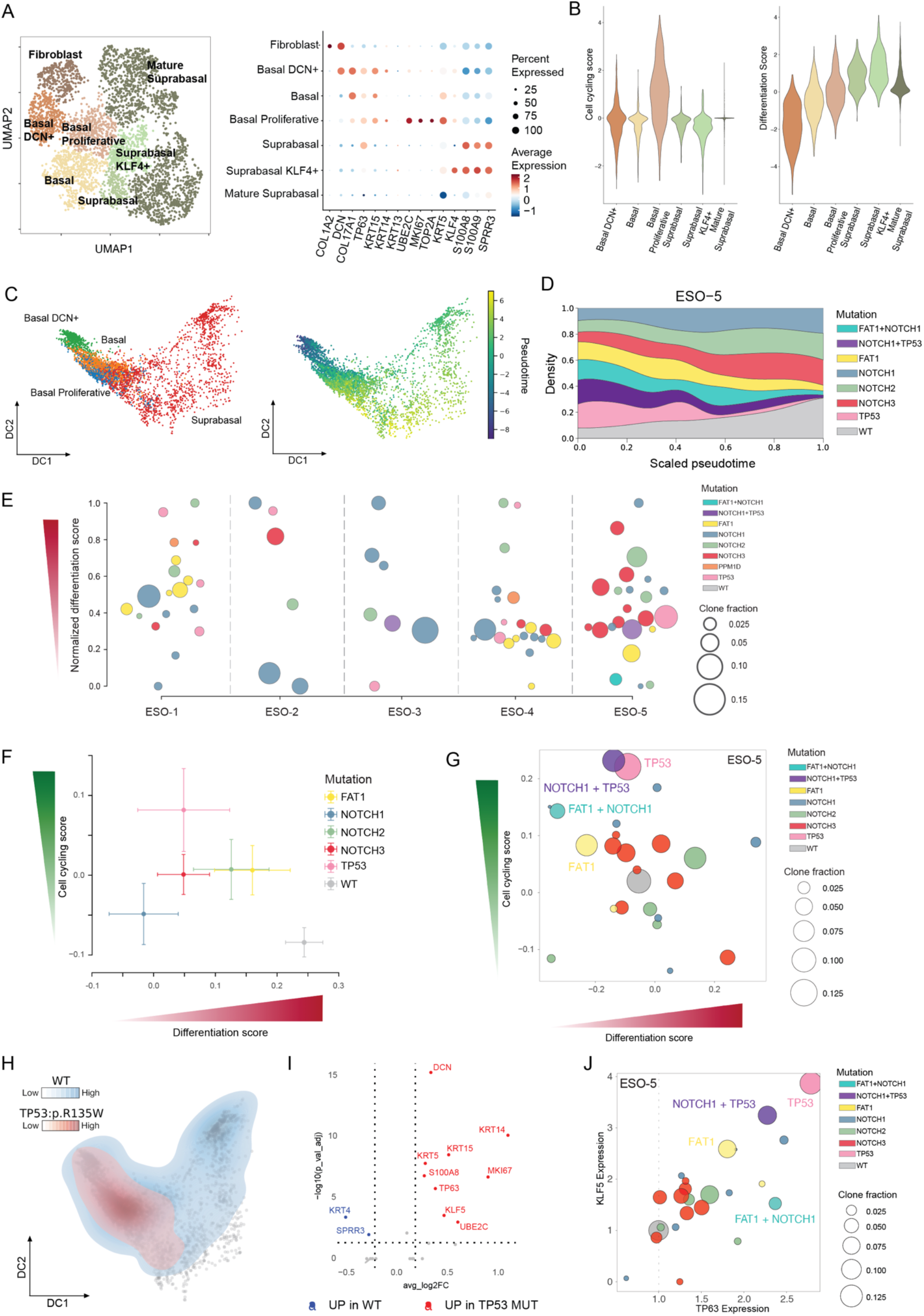
Esophageal epithelium is colonized by diverse somatic mutant clones with phenotypic biases. **A**. (Left) Uniform manifold approximation and projection (UMAP) of single cells from all samples using RNA expression, colored by annotated cell types. (Right) Dot plot displaying cell type marker genes (x-axis) used for cell type annotation (y-axis). Dot size represents the percentage of cells expressing the marker gene, and color scale indicates the mean expression level (centered log ratio) within each cell type. **B**. Violin plots comparing cell cycling scores (left) and differentiation scores (right) across assigned cell types. Cell cycle and differentiation scores were calculated as gene-module scores based on cycling and differentiation gene sets expression from the RNA panel. **C**. Diffusion map of epithelial single cells merged from ESO-1, ESO-2, ESO-3, and ESO-5, annotated by cell types (top) and overlaid with trajectory scores (bottom). The trajectory scores represent the differentiation stage of each cell along the inferred pseudotime trajectory. Fibroblasts from **A** were excluded from this analysis. **D**. Clonal composition across the pseudotime diffusion map representing the differentiation stage for the ESO-5 sample. Clone fractions are summed by driver gene mutation and relative abundance plotted against pseudotime quantiles representing differentiation trajectory. **E**. Mean differentiation scores of cells assigned to clones across all donor samples. Dot size represents the clone fraction and color indicates the mutant driver gene. The differentiation scores are min-max scaled within each sample to allow for comparison across samples. **F**. Mean differentiation and cycling scores of cells assigned to clones with single driver mutation, aggregated by mutant driver genes across all samples. Each point represents aggregation of cells with mutation in a specific driver gene, with error bars indicating the standard error of the mean across cells (SEM). **G**. Clones from the ESO-5 sample projected onto the differentiation and cell cycling score axes (mean score of cells assigned to clones). Each dot represents a clone, with the dot size proportional to the clone fraction and color indicating the mutant driver gene. WT clone fraction is fixed at 0.1 for visualization purposes. True WT proportions represented in Fig. 1H. **H**. Diffusion map from **C** overlaid with kernel density estimates over those dimensions for cells with *TP53*:p.R135W mutation. **I**. Volcano plot comparing differentially expressed transcripts between *TP53* mutant cells and wild-type cells. Horizontal dotted line represents FDR < 0.05; vertical dotted lines represent average log2 fold change (log2FC) > 0.15. **J**. Clones from the ESO-5 sample projected according to their mean expression of *KLF5* and *TP63*. Dot size represents the clone fraction, and color indicates the mutant driver gene. WT cells are fixed at clone fraction of 0.1 and used to visualize as comparison to clone scores.

Transgenic mouse models have shown that widespread *Notch1* loss has minimal effects on epithelial structure and cell dynamics, but increases clonal fitness^14^. The dividing progenitor cells in the basal compartment must generate either two progenitor cells, two differentiating cells, or one of each^15^. Wild-type tissue retains equal proportions of both progenitor and differentiated cells to maintain homeostasis, but mutations that bias differentiation to progenitor-like cells can drive clonal expansion. Furthermore, lineage tracing experiments in murine models have demonstrated that heterozygous *TP53* mutants have similar effects in inducing higher rates of progenitor production while maintaining epithelium structure^32,33^. To assess the impact of mutations on differentiation status of somatic clones in our PNE samples, we compared differentiation scores across wild-type cells and clonal populations (**Fig. 2E**, Methods). This analysis showed a wide distribution of differentiation biases across mutant clones compared to wild-type cells (**Fig. 2E**). For instance, 7/11 clones with a *TP53* mutation and 18/28 clones with a N*OTCH1* mutation had lower differentiation scores compared to wild-type cells, indicating that the majority of clones with these driver gene mutations exhibited a bias towards earlier differentiation states. Notably, the largest clone in every sample was represented within these early cell state-biased clones (**Fig. 2E, Supplementary Fig. 8)**. Moreover, double-mutant clones also showed increased bias towards early differentiation, despite different combinations of mutations **(Fig. 2E, Supplementary Fig. 8)**. Finally, when clones were stratified by the mutant driver gene, *TP53, NOTCH1*, and *NOTCH3* mutants were most biased towards early differentiation states (**Fig. 2F**). These findings demonstrate that similar to animal models, in human PNE these driver gene mutations are associated with a differentiation bias that may underlie clonal expansions. Furthermore, our phenotypic mapping helps explain previous findings that the acquisition of additional driver mutations induces clones that span larger spatial areas^12^, as all double mutant clones were biased towards earlier differentiation states.

While differentiation biases may contribute to clonal expansions, driver mutations may also directly impact cell proliferation as an alternative route to enhanced fitness. We therefore compared the differentiation state and cell cycling activity of single-mutant clones **(Fig. 2F)**. Notably, while *TP53* mutants showed both early differentiation and higher cell cycling activity, *NOTCH1* mutants only showed early differentiation bias **(Fig. 2F)**. For the double mutant clones (detected in two samples), we observed bias towards early differentiation or increased proliferation. Specifically, in ESO-5, a *FAT1+NOTCH1* mutant clone and its parent *FAT1* mutant clone were both highly skewed towards early differentiation state and increased proliferation score. Another *TP53+NOTCH1* mutant clone and its parental *TP53* mutant clone had the highest cell cycling score compared to wild-type and other mutant clones in this sample (**Fig. 2G**). In ESO-3, a *TP53+NOTCH1* mutant clone and its parental *TP53* mutant clone both were highly biased towards early differentiation (**Supplementary Fig. 8**). These data suggest that driver gene mutations may promote clonal expansion through mechanisms that stall differentiation or increase proliferation, and in some cases through the combination of both.

We observed that ESO-5 contained cells with a *TP53*:p.R135W mutation that are phenotypically biased along the epithelial differentiation **(Fig. 2H)**. Although the increased fitness of *TP53* mutant cells has been documented, especially in the context of ESCC development^33^, the molecular basis in the context of PNE is not well understood. When compared to wild-type cells in the same donor, *TP53*-mutant cells have overexpression of basal cell markers (*KRT14, KRT15*), cycling genes (*UBE2C, MKI67*), and keratinocyte differentiation regulators (*TP63, KLF5*) (**Fig. 2I**). Notably, p53 normally interacts with regulators of differentiation, including p63 and members of the *KLF* family (e.g., *KLF4 and KLF5*) that are commonly dysregulated in esophageal cancers^34,35^. As such, disrupted p63 and KLF signaling may underlie the bias towards earlier differentiation states in these *TP53*-mutant clones. Indeed, in our oldest donor ESO-5, cells in the large *TP53*-mutant clone (along with its subclone harboring additional *NOTCH1* mutation) had overexpression of *TP63* and *KLF5* compared to wild-type cells (**Fig. 2J**; see **Supplementary Fig. 9** for *TP63* and *KLF5* expression profiles for other donors). Previously, it was shown that loss of p53 in epithelial layers of the esophagus in mice increased p63 levels, allowing the cells to activate p21 in lieu of p53 to maintain tissue integrity^36^. In addition, *TP63* expression is a known marker of stem and progenitor basal cells^37^, and it has been hypothesized to act as a switch between proliferating basal to differentiating suprabasal cells^36^. These reports suggest that the increase in *TP63* expression that we observe in our *TP53*-mutant clones could allow these cells to respond to DNA stress, maintain early progenitor state and stall differentiation. In turn, these phenotypes could increase the fitness of *TP53*-mutant cells over wild-type cells. Together, we demonstrate the linkage of phenotypic readouts with a cell’s genotype is required to uncover how stalled differentiation and enhanced proliferation can enable mutant cells to gain fitness advantages in PNE without drastically disrupting tissue composition of cell dynamics.

## Discussion

Clonal mosaicism in malignancy has long been recognized as a critical feature that allows cancer cells to increase fitness and develop resistance mechanisms through evolutionary selection. Recent work has demonstrated the ubiquity of CM also across normal aging tissues and in non-malignant disease contexts. For example, sequencing of liver microdissections identified positively-selected driver mutations in metabolic genes, including *FOXO1, GPAM* and *CIDEB*, in patients with non-alcoholic fatty liver disease^10^. In inflammatory bowel disease, somatic mutations have been identified in the colonic epithelium, with driver mutations reported in key inflammatory signaling genes, including those in the IL17 pathway^7,8^, further underscoring how disease environments can select for functionally consequential somatic mutations. Notably, even non-diseased tissues harbor high levels of clonal expansions; phenotypically normal sun-exposed skin is overtaken by clones that acquired mutations in cancer-related genes^31^. Similarly, the phenotypically normal esophagus was reported to contain nearly ubiquitous clonal expansions with age, further increasing with tobacco and alcohol exposure^4,12^. Collectively, these findings show that the complex process of human somatic evolution is influenced by aging, environment and disease state, suggesting an intricate interplay between somatic mutational genotype and cellular phenotype.

Bulk genomic characterization of solid tissue CM such as PNE has revealed the extent of clonal expansions, but not at resolutions required to identify all incidences of co-occurrence of mutations in clones. While mutations with large VAFs allow clonal phasing through inference methods (e.g., pigeonhole principle that requires the sum of two corrected VAFs > 1.0 to determine clonal relationship^4,5,31^), bulk studies require precise optimization of tissue size to achieve a balance of obtaining sufficient DNA for sequencing and to maintain a relatively large clone size within the sampled tissue. For example, in bulk deep-targeted sequencing of PNE, only 25 out of 844 microdissections had large enough clones to determine co-occurring driver mutations in the same cells and the majority of variants detected had mutant cell fractions < 0.2 of the sampling area^4,12^. A single-cell perspective can identify whether mutations are nested within the same clone, or are present in sibling clones. For example, scDNAseq methods have revealed high resolution clonal architecture in cancers (AML, CLL, breast cancer), but have not been applied to solid tissue CM.

Using scG2P, we analyzed normal esophagus biopsies from aged donors to delineate clonal structure and mutation complexity. By targeting the six most highly mutated driver genes, we detected a driver mutation in more than half of cells assayed, aligning with previous discoveries that driver mutants colonize the esophageal epithelium. Our ability to resolve clonal hierarchies shed light on previous mutational profiling, demonstrating that driver mutations induce clones of varying sizes. These data address a key question in the field of CM. While the same driver gene mutations are also observed in cancer, malignant clones typically harbor multiple drivers^38,39^. In contrast, in CM of phenotypically normal tissue, we show through our single-cell DNA sequencing that clones harbor mostly single driver mutations, with only rare instances of two driver mutations in the same clone.

By adding the capture of targeted mRNA markers with genotyping in scG2P, we demonstrate the first genotype-to-phenotype mapping of solid tissue CM. Through a cell line mixing study, we showed that scG2P can jointly capture genotype and phenotype at high accuracy. The application of scG2P to PNE samples allowed the measurement of RNA markers of differentiation and cell cycle, and linked them to driver mutation identity, number of mutations and clone size. We inferred clonal fitness based on clone fraction, and found that clones with the highest fitness were biased towards early differentiation phenotypes. In *NOTCH1*, the most commonly mutated driver gene in PNE, we observed stunted differentiation compared to wild-type cells within the same sample. Importantly, this observation was obtained from primary human samples, using wild-type cells as the ideal control as they originate from the same individual and micro-environment. This finding aligns with murine models that show that *Notch1* mutation leads to clonal outgrowth through increased symmetric progenitor divisions at the expense of contribution to differentiated cells^14^. Of note, *NOTCH1* mutations are observed less frequently in cancer than in PNE^12^, suggesting a protective role for *NOTCH1* loss in carcinogenesis. This is consistent with scG2P data showing that *NOTCH1* mutants do not drive increased proliferation in the PNE. In contrast, *TP53* mutants showed both differentiation biases and increased proliferation as drivers of clonal outgrowth. We further link *TP53* mutants to increased *TP63* expression, suggesting a pathway to increased clonal fitness, as elevated levels of p63 have also been observed both in esophageal tumors as well as adjacent normal tissues, suggesting its role as an early event in carcinogenesis by field effect^40^. Finally, rare double mutant clones showed the highest phenotypic differences compared to wild-type cells, with both greater proliferation and more pronounced differentiation biases. These data support the model in which acquisition of multiple drivers is required for a stepwise malignant transformation process, and suggest that tissue integrity of the esophagus is maintained despite the overwhelming clonal driver gene colonization due to the fact that only rarely clones harbor more than one driver.

While our work presents high-resolution multimodal sequencing of the human CM, we note several limitations. First, our analysis likely underestimates the true extent of clonal diversity and tissue colonization due to the technical challenges associated with high throughput single-cell mutational profiling, as increasing the number of targeted genes increases primer-primer interactions and decreases capture efficiency. It is also technically challenging to effectively capture regions with high G-C content, illustrated by the large numbers of *NOTCH1* amplicons with decreased capture efficiency, as compared to the efficiency across *TP53* and *FAT1*. These design challenges likely lead to underestimation of the frequency of single and double driver mutant clones. However, this is likely a modest effect, as our sensitivity analysis confirmed that we observe similar clonal proportions when increasing coverage stringency. Second, we note that there is a high prevalence of copy-neutral loss of heterozygosity (CN-LOH) events, such as *NOTCH1* loss, that have been observed in aged PNE. scG2P was only able to detect a portion of the expected copy number changes in our cell line mixing study, and we hypothesize an extension of the panel to include genomic loci that include germline SNPs in future panels can be used to more accurately detect these events in tissues. Last, our mRNA panel is limited in size. Future technology development will aim to increase the breadth of the genome and transcriptome capture at single-cell resolution, while retaining the advantages of throughput afforded by droplet microfluidic empowered single-cell genomics that can profile thousands of cells in a single experiment.

In summary, clonal diversification is a hallmark of human somatic tissue, in both malignant and nonmalignant contexts. However, the ability to determine how clones outgrow requires the ability to link genotype to phenotype in primary human samples. scG2P provides a novel platform to interrogate clonal diversification and the resulting cellular differentiation biases at the throughput necessary to address human clonal complexity. This technology specifically empowers the study of human clonal mosaicism in solid tissues by profiling multiple variants along driver genes together with analysis of cell state, cell cycle status and differentiation stage. As such, our technology addressed a critical gap in single-cell genotype-phenotype mapping, where technologies have been limited in throughput^17,41^ and in targeting isolated hotspots^19,20,42^. Moreover, scG2P addresses specific challenges in human solid tissue mosaicism by enabling the study of nuclei from archival tissues, with the throughput and multiplexing capability required for the study of clonally complex samples. This framework is poised to advance our understanding of CM as one of the most exciting frontiers in human genetics, with major opportunities for discovery related to cancer^9,13^, aging^4,12,44^, non-malignant chronic disease^7–11^, and pre-cancer states^45–47^.

## Methods

### Esophageal biopsy collection

We enrolled patients who underwent therapeutic or diagnostic endoscopy for upper gastrointestinal symptoms at Kyoto University Hospital. Informed consent was obtained from all participants. Characteristics of these patients and healthy individuals are summarized in Supplementary Information. A history of heavy alcohol drinking (HIGH indicated by ≥396 g alcohol per week) or tobacco smoking (HIGH indicated by ≥30 pack-years) was reported in these individuals, who were considered to have positive lifestyle ESCC risks (high-risk individuals). Biomaterials, including esophageal tissues, were newly collected from 11 individuals using endoscopic biopsy, according to procedures approved by the Internal Review Board at Kyoto University (G0645).

### Tissue processing

#### Single cell dissociation

Tissue biopsies were rinsed in ice cold 1X PBS and kept in ice cold Keratinocyte Serum Free Medium (KSFM, ThermoFisher) immediately after collection until processing. Tissues were placed with 500 μL ice cold 1X PBS and minced with surgical scissors. 500 μL of 0.25% trypsin (ThermoFisher) were added to the sample and incubated at 37 °C, with vortexing every 10 minutes. Reaction was stopped by adding 1 mL of KSFM with 10% FBS and strained through a 70 μm strainer. Cells were collected by centrifugation at 300 g for 5 minutes and washed with cell staining buffer (Biolegend). Cells were incubated with FcX (Biolegend) for 10 minutes at 4 °C and then stained with FITC - anti-human EPCAM (BioLegend CAT #32404) and PE - anti-CD45 (BioLegend, CAT # 368510) for 30 minutes at 4 °C before sorting for CD45-negative and EPCAM-positive cells.

#### Single nuclei dissociation

Tissue biopsies that were OCT embedded were trimmed of excess OCT, and then loaded on Singulator along with final concentration of 1.0 U RNAse inhibitor (Protector, Sigma) and 1mM DTT using standard nuclei extraction protocol. Sucrose (final concentration 250 mM) was added to nuclei output to mix and spun down at 500 g, 4 °C, for 5 minutes. Nuclei were washed with nuclei wash buffer (1X DPBS, 1.0 U RNAse inhibitor, 1mM DTT, 1% BSA) twice and filtered through a 40 μm strainer. Nuclei were stained with DAPI and counted on a Countess.

### 10x Chromium Single cell RNAseq / 10x Visium Spatial Gene Expression

Single-cell RNAseq was carried out according to the manufacturer’s standard protocols using 10x Chromium 3’ Single Cell Gene Expression protocol. For 10x Visium assays, tissue biopsies were OCT embedded and stored. On the day of analysis, extra OCT was trimmed and tissue cryosectioned to 10 μm thickness and placed on Visium Spatial Gene Expression slides. Spatial transcriptomics was then carried out according to the manufacturer’s standard protocols. Fastqs generated from Illumina sequencing was input to cellranger (v7.1.0, Chromium scRNAseq) and spaceranger (v1.3.1, Visum spatial transcriptomics) with standard parameters aligning to hg38.

Cellranger and spaceranger outputs were exported to the R package Seurat^48^ (v4.0.0) for downstream analysis. Standard pre-processing workflow was used to QC and select cells (number of genes > 200, and % mitochondrial reads < 5). Log normalization was applied to the data, with the top 2,000 variable features used for principal component analysis. The top 15 principal components were used for graph-based clustering and non-linear dimensional reduction to uniform manifold approximation projection (UMAP). The reference esophageal scRNAseq dataset from Madissoon et al.^29^ was downloaded from CellxGene, and a Jacard similarity score was calculated between clusters and cell type annotations provided by authors. Cluster annotation was performed manually by examining Jaccard similarity scores. The top differentially expressed genes were determined by Seurat’s FindMarkers function and compared to marker genes provided in Madissoon et al.^29^

### Cell lines

KYSE-410 (Sigma-Aldrich CAT # 94072023-1VL), KYSE-270 (Sigma-Aldrich CAT #94072021-1VL), and HCT-116 (ATCC CAT #CCL-247) were cultured according to manufacturer instructions before loading onto Tapestri platforms. Cell lines in culture were screened biweekly for mycoplasma contamination using the MycoAlert PLUS Mycoplasma Detection Kit (Lonza, 801 #LT07-703).

Whole-exome sequencing (WES) data for each cell line were downloaded from DepMap portal (depmap.org, DepMap Public 22Q1). CCLE_mutations were extracted to filter for variants intersecting with our scG2P mutation panel.

### scG2P RNA/DNA

#### DNA panel design

Our DNA panel was designed using the Mission Bio Tapestri Designer by inputting all mutations detected in Yokoyama et al.^12^ to generate amplicon primers. The Mission Bio panels consist of amplicons ranging in size from 175 to 275 bp generated using primers of 18–35 bp in length. Amplicons can have decreased targeting performance in regions with high %GC content or that are highly repetitive. The final DNA panel size was composed of 118 DNA amplicons covering 7 driver genes. No variants were detected in *ZFP36L2* (1 amplicon) and were not included in further analysis.

#### RNA panel design

A panel of 86 targets was originally designed to capture esophagus-specific mRNA targets. For this design, the most frequent exon-exon junction was selected for each gene. The transcript of interest was confirmed to contain the exon-exon junction targeted and the amplicon was designed to cross these exons, allowing for differentiation from cDNA amplification and gDNA amplification by the presence of the intron sequence in the amplicon. Targets were removed if they could not include an exon-exon junction.

The primers were designed as 12-30 nt with ∼20 nt preferable, within a GC range of 50-60%, and for amplicon lengths of 200-400 bp. A single set of capture primers was selected for each gene target. These primers were confirmed not to contain any common SNP sites at the 5 bases located at the 3’ end of the primer. The Gibbs free energy was also calculated and any RNA reverse primers with a low ΔG (ΔG < -6 kcal/mol using the OligoAnalyzer tool on https://www.idtdna.com/pages/tools/oligoanalyzer) were redesigned. Additional checks were performed to minimize primer interactions. To assess all primers (DNA and RNA) for interactions, primers were checked to ensure that the 8 bases of the 3’ end were not an exact match with the 8 bases of the 3’ end of any other primer. A more stringent primer interaction check for the primers used for reverse transcription were performed only with the RNA reverse primers. The Multiple Primer Analyzer tool (Thermofisher) was used to confirm that the 5 bases at the 3’ end did not have an exact complementary match to any other RNA reverse primer, including the adaptor sequence. Further targets were excluded if they were not amplified or non-specifically amplified in the cell line mixing experiment.

Primer PCR handles were added to the RNA primers. The forward primers had 5’-GTACTCGCAGTAGTC-3’ appended to the 5’ end of the gene-specific primer and the reverse primers had 5’-GTGATACACGACTATGAGCGCTA-3’ appended to the 5’ end of the gene-specific primer.

#### Tapestri RNA+DNA protocol

Single cells or nuclei were counted using Trypan exclusion using a Countess 3 and resuspended in nuclei wash buffer (1X DPBS, 1.0 U Protector RNAse inhibitor, 1mM DTT, 1% BSA) at concentration between 1,000-3,000 cells per μL to reach a minimum volume of 35 μL. The Mission Bio v2 chemistry was used for this protocol. Step one of the Mission Bio Tapestri protocol was performed by adding in the cell suspension and encapsulation oil v2 (Mission Bio) to the Tapestri cartridge along with a mastermix containing RT and cell lysis reagents. This mastermix contained final concentrations of 10 U/μL of SuperScript IV (Thermo #18091050), 5 mM DTT (Thermo Fisher #18091050), 0.5 mM each dNTP (Thermo Fisher #18091050), 1X 5X buffer (Thermo Fisher #18091050), 2 U/μL Rnase inhibitor (Thermo Fisher #18091050), 1% NP-40 Surfact-Amps™ Detergent Solution (Thermo Fisher #85124), 0.04 mg/mL Proteinase K (Sigma #3115828001) and 14.6 μM RNA reverse primer pool. The Proteinase K was diluted in nuclease free water prior to adding it to the mastermix and was added immediately before loading on the Tapestri cartridge to avoid degradation of the enzymes.

| Reagent | Volume (μL) |
| --- | --- |
| nuclease-free water | 42.8 |
| 5X buffer | 20 |
| NP-40 Surfact-Amps Detergent Solution (10%) | 10 |
| RNA reverse primer pool (200 μM total) | 5 |
| DTT (0.1 M) | 5 |
| dNTPs (10 mM each) | 5 |
| RNase Inhibitor (40U/μL) | 5 |
| SuperScript IV enzyme (200U/μL) | 5 |
| Proteinase K (1.9 mg/mL in nuclease-free water) | 2.2 |

Encapsulation protocol (Mission Bio) was run following manufacturer’s instructions to generate the first emulsion. A gel loading tip was used to remove excess encapsulation oil from the emulsion tube. The lysis and digestion protocol was run on a preheated thermocycler with the following thermocycler protocol:

| Step | Temperature ($^{\circ}$C) | Time |
| --- | --- | --- |
| 1 | 50 $^{\circ}$C | 60 min |
| 2 | 80 $^{\circ}$C | 10 min |
| 3 | 4 °C | HOLD |

Emulsions were loaded onto Tapestri cartridge with Barcoding Mix (293.8 μLBarcoding Master Mix v2 with final concentrations of 0.21 μM DNA forward primer pool, 2.1 μM DNA reverse primer pool, 0.42 μM RNA forward primer pool), Barcoding beads, and Barcoding oil v2 (Mission Bio). Barcoding program was run to generate a second emulsion and UV light was used to cleave off barcode-containing forward primers from Barcoding beads prior to PCR amplification. A gel loading tip was used to remove excess barcoding oil from the emulsion tubes.

| Reagent | Volume (μL) |
| --- | --- |
| Barcoding MM v2 | 293.8 |
| DNA forward primer pool | 2.5 |
| DNA reverse primer pool | 3.1 |
| RNA forward primer pool (200 μM total) | 0.6 |

Emulsions were loaded onto the Tapestri cartridge with Barcoding Mix (Barcoding Master Mix, DNA forward primer pool, DNA reverse primer pool, RNA forward primer pool), barcoding beads, and barcoding oil (Mission Bio). Barcoding program was run to generate a second emulsion and UV light was used to cleave off barcode-containing forward primers from barcoding beads prior to PCR amplification. A gel loading tip was used to remove excess barcoding oil from the emulsion tube.

Targeted PCR was performed using the following thermocycler protocol:

| Step | Ramp (°C/second) | Temperature (°C) | Time | Cycles |
| --- | --- | --- | --- | --- |
| 1 | 4 | 98 | 6 min |  |
| 2 | 1 | 95 | 30 sec | 10 |
| 3 |  | 72 | 10 sec |  |
| 4 |  | 61 | 6 min |  |
| 5 |  | 72 | 20 sec |  |
| 6 | 1 | 95 | 30 sec | 12 |
| 7 |  | 72 | 10 sec |  |
| 8 |  | 51 | 4.5 min |  |
| 9 |  | 72 | 20 sec |  |
| 10 | 4 | 72 | 2 min |  |
| 11 |  | 4 | HOLD |  |

After targeted PCR, 15 μL of an extraction agent (Mission Bio) was added to samples to break emulsion. DNA clean up buffer and clean up enzyme were added to sample and incubated at 37 °C for 60 minutes at 350 RPM. Two AMPure XP library clean-up steps were performed at 0.72X.

The RNA PCR product isolation was performed by washing M-270 Streptavidin beads (ThermoFisher) in Binding and Washing Buffer (10M Tris-HCL, 1 mM EDTA, 2M NaCl). An RNA Biotin Oligo (/5BioTinTEG/GTGATACACGACTATGAGCGCTA/3C6/) was added to the PCR product at a final concentration of 0.7 μM. The PCR product with the RNA Biotin Oligo was incubated at 99 °C for 5 minutes and transferred immediately to ice for 5 minutes. The Streptavidin beads were added and incubated at room temperature for 30 minutes before magnetic separation of the DNA and RNA fractions. A 0.72X AMPure cleanup was then performed on the DNA fraction.

Both fractions undergo library amplification using the entire product using Library Mix (Mission Bio). The DNA PCR product was amplified with the Mission Bio kit indexes (Mission Bio) and the RNA PCR product was amplified with custom indices using the following thermocycler protocol:

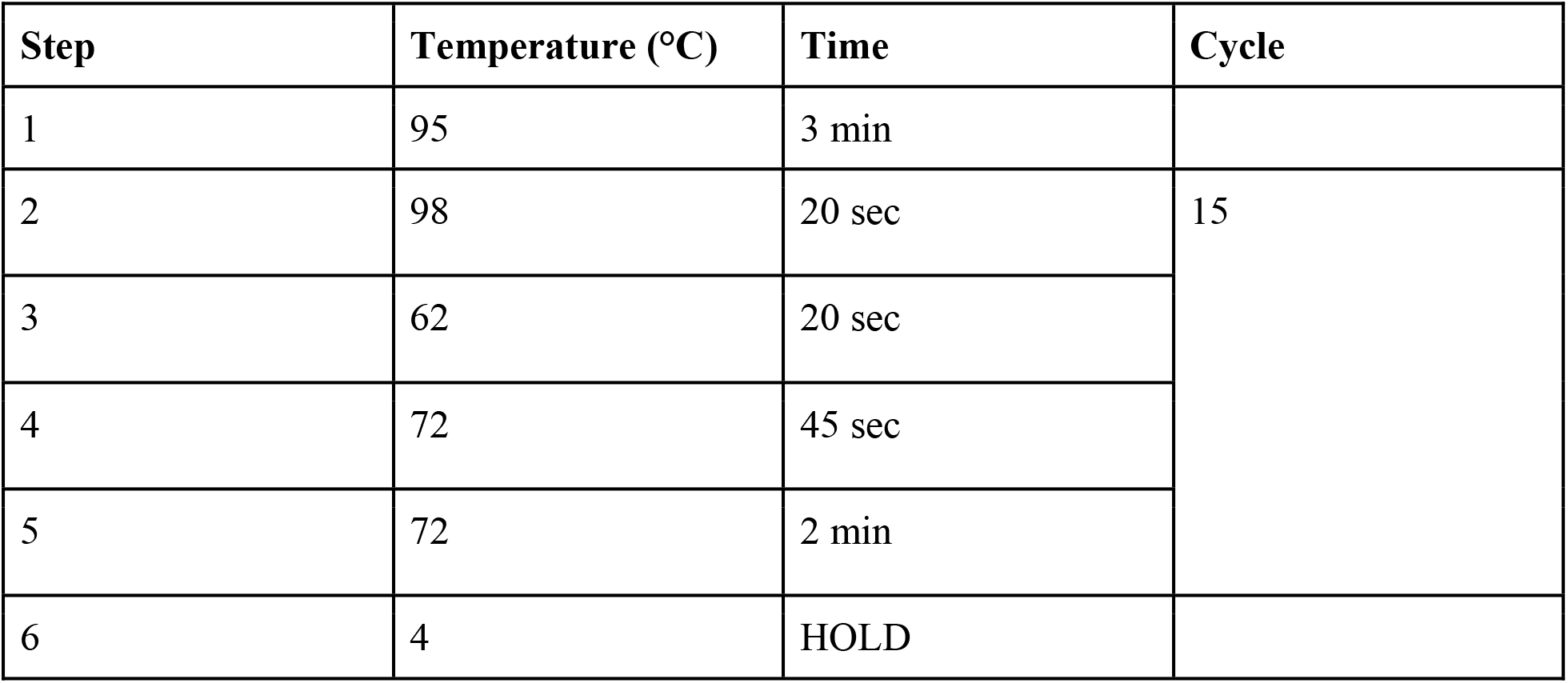

All libraries were sequenced with paired-end 150 bp on Illumina Novaseq 6000 at the Weill Cornell Medicine Genomics Core.

### Data processing

Fastqs for DNA libraries generated were processed using the Tapestri pipeline (Mission Bio) for adapter trimming, barcode correction, sequence alignment, cell calling, and variant calling. RNA libraries were processed for adapter trimming and sequence alignment with STAR.

#### Cell calling

Cell finding from the DNA library was performed using a “Correlation UMAP” algorithm. Given a barcode by amplicon read count matrix, two metric matrices were derived, normalized read counts and correlation-coverage. Normalized read counts were generated by first normalizing each amplicon read count to its mean plus one across all barcodes, then normalizing these values to the median read counts of the top 10% of all barcodes plus a small constant. Correlation-coverage was derived from calculating the log10 of the mean coverage for the barcode across all amplicons and the $r^2$ of the per-amplicon coverage for the barcode with that of the experiment as a whole. These two feature sets were then combined and scaled to have equal weighting under L1-distance. UMAP was then performed on this combined feature set. The resulting UMAP was then clustered with HDBSCAN and the resulting clusters were used to identify the cell cluster based on quality control metrics.

#### Variant calling

Mission Bio’s Tapestri-2.0.1 pipeline employs a barcode decoding step to demultiplex the sequence data, followed by BWA mapping of the sequences. Variant calling is performed on a per-cell basis with GATK HaplotypeCaller (v4.1.7.0). The set of GVCFs are then merged and converted to a MissionBio object for further analysis. Filtering of the variants was performed with MissionBio’s Mosaic v2.1 package using the following settings: variants must have a minimum depth of 10 reads, minimum genotype quality of 30, and be called in a minimum of 50% of the cells.

### Clonal reconstruction and genotyping completeness

1) We implemented an unsupervised iterative soft-clustering algorithm to assign cells to cluster genotype states that maximizes each cell’s membership vector. Sites where genotypes were missing in a cell had their weight redistributed evenly to all other valid sites in our distance metric. 2) We implemented a reinforcement learning clonal trajectory model based on Miles et al.^21^ to reconstruct clonal structure. Briefly, a reward matrix was constructed with rewards proportional to genotype state prevalence observed in the sample. The action space was restricted to those that define transitions where a single site is mutated and infinite sites is satisfied. A Q-learning agent was trained using an epsilon-greedy with experience replay algorithm to learn the values of each genotype state transition. The resulting Q-matrix was then transformed into a genotype state graph with states are nodes and mutational events as edges weighted by their Q value. The optimal path to each genotype state observed in the sample was extracted to produce the resulting optimal clonal trajectory. We filtered out clones that were not reproducible with bootstrapping methods^21^. Additionally, to examine whether clone sizes were correlated with genotyping completeness (**Supplementary Fig. 6C**), we thresholded for cells with >50% and >80% genotyping across all variants and observed no clone size biases based on genotyping completeness. We performed this analysis specifically for clones with nested double mutations where the single mutant clone is also detectable in the same sample (**Supplementary Fig. 6E**). To account for the possibility of allelic dropout at the second site, we used high levels of stringent filtering to confidently detect single and nested double mutant clones, and verified that the ratios of both were consistent between low and high genotyping completeness thresholds.

### RNA Sequencing Reads Filtering

To mitigate potential gDNA contamination in the scRNA panel, we developed a computational pipeline, *PRIMR*, that selectively identified reads exhibiting exon-exon junction. First, the BAM file outputted by STAR (v2.7.0f) was filtered to only include reads with no more than 5 bases aligned to an intronic region. This BAM file was then partitioned into amplicon-specific BAM files, where each file corresponded to a distinct amplicon (e.g., a gene from the panel). These BAM files were then sorted based on the read names (QNAMES). Next, a paired-end bed file-like file was generated, with each row representing a read and its mate (R1/R2) from the same fragment. The start and end positions of the reads were updated using the CIGAR string information, accounting for soft-clipped bases in the position calculations. This approach allowed for the identification of reads derived from gDNA, which mapped to the intronic region but had bases soft-clipped to exclusively align to the exonic region by the STAR splice-aware aligner. By utilizing the CIGAR-corrected positions, fragments mapping to intronic regions were filtered out, retaining only those fragments that exhibited exon-exon junctions in at least one of their reads (e.g., read 1 (R1) or read 2 (R2)). These fragments were subsequently employed to compute the corrected RNA counts matrix (**Supplementary Fig. 4**).

### Cell Line Mixing Data Processing and Analysis

#### DNA Data Processing and Analysis

Raw DNA sequencing data were extracted from the h5 file generated by the Mission Bio Mosaic pipeline. To reduce the dimensionality of the DNA VAF data, the UMAP algorithm implemented in the RunUMAP function of the Seurat package (v5.0.0) was used. The set of identified features was provided as input, and parameters seed.use and min.dist were set to 123 and 0.01, respectively. This allowed us to visualize the high-dimensional data in a two-dimensional space. Subsequently, k-means clustering with k=3 was performed to identify the three distinct cell lines based on the allele frequencies of SNVs. Clusters were labeled using the SNVs that were unique to each cell-line, as shown in **Fig. 1C**.

### RNA Data Processing and Analysis

For RNA data processing, the *PRIMR* R package (https://github.com/Theob0t/PRIMR), developed by our team, was employed to filter the raw sequencing reads to retain only exon-exon junction reads (method detailed below). Droplets identified as cells by the Tapestri cell caller and expressing more than 100 RNA reads were retained. To address potential multiplets and exclude remaining empty droplets, barcodes expressing a number of unique features beyond the 95th percentile or below the 5th percentile of cells were removed. The processed RNA counts matrix was normalized using a Centered-Log-Ratio (CLR) transformation across cells as defined in the Seurat package. Principal Component Analysis (PCA) was then performed on all features to reduce dimensionality. Further dimensionality reduction was achieved using the RunUMAP function with the first 5 principal components, and parameters seed.use and min.dist set to 123 and 0.01, respectively. To identify clusters of cells, the Louvain algorithm was employed, calculating k-nearest neighbors and constructing the shared nearest neighbor (SNN) graph using the FindNeighbors function. The FindClusters function with a resolution parameter of 0.15 was used to determine the clusters.

#### Visualization and Cell Line Assignment Accuracy

A plot of the UMAP space demonstrated clear separation of the three cell lines, marked by expression of their respective markers *KRT5, KRT7*, and *KRT23*. The accuracy of the cell line assignment was calculated by computing the ratio of cells assigned to the same cell lines using both the RNA and DNA panels and the total number of cells.

#### Correlation Analysis between scG2P scRNA Data and CCLE Bulk Reference Datasets

Bulk RNA sequencing data for HCT116, KYSE270, and KYSE410 cell lines were obtained from the CCLE database and the reported log2(TPM+1) values were used for the analysis. scG2P single-cell RNA sequencing data were pseudo-bulked based on the DNA VAF clustering labels, using the average of the Centered Log-Ratio values normalized across features. Correlations were computed using the Spearman correlation coefficient and visualized per cell line using scatter plots.

#### Correlation Analysis between scG2P Single-Cell Data and Drop-seq Dataset

HCT116 single-cell data sequenced from the Park et al.^49^ study using the Drop-seq protocol were downloaded from the GEO database (GSE149224). Data filtering was performed using the GSE149224_meta.information.csv file to retain only HCT116 cells with a dose equal to 0. scG2P single-cell RNA sequencing data were pseudo-bulked based on the DNA VAF clustering labels, using the average of the Centered Log-Ratio values normalized across features. Correlations were computed using the Spearman correlation coefficient and visualized per cell line using scatter plots.

### Patient Samples Data Processing and Analysis

#### RNA Data Processing and Analysis

Raw RNA data for each patient were processed by the *PRIMR* pipeline to select only for fragments exhibiting an exon-exon junction. Corrected RNA counts matrices were merged and droplets identified as cells by the Tapestri cell caller were retained. Cells expressing fewer than 50 RNA reads were removed from the analysis. The RNA counts matrix was normalized using a Centered-Log-Ratio (CLR) transformation across cells as defined in the Seurat package. To account for variations in RNA sequencing depth, the ScaleData Seurat function was applied with the vars.to.regress parameter set to nCount_RNA; both the center and do.scale parameters were set to False. Dimensionality reduction was performed using the RunUMAP function on the normalized matrix, with parameters seed.use and min.dist set to 123 and 0.5. FindNeighbors Seurat function was used to compute the SNN graph. Subsequently, the FindClusters function was employed to cluster cells using the Louvain algorithm, with a resolution parameter set to 0.15.

#### Cell Cycling and Differentiation Module Scores

Cell Cycling and Differentiation module scores were calculated using the AddModuleScore Seurat function. Briefly, the function calculates the average expression levels of each module at the single cell level, subtracted by the aggregated expression of control feature sets. All analyzed features are binned based on averaged expression, and the control features are randomly selected from each bin. For this analysis, we used 10 control features per analyzed feature and 5 bins of aggregate expression levels for all analyzed features.

#### Diffusion map and trajectory inference

Fibroblasts were removed from gene counts matrix and a diffusion map was generated from the neighbors graph of the scRNA data as implemented in scanpy^50^ (v1.9.0) and diffusion pseudotime calculated along the diffusion components while rooted in our basal cell cluster. Clonal abundance fish plots were generated by calculating density of mutant cells with driver gene mutations over pseudotime quantiles, representing trajectory of differentiation from basal to differentiated. ESO-4 was not used for diffusion map construction due to the lower number of features detected compared to other samples.

### Plotting

All schematics were generated in Biorender. Plots were generated with ggplot2 (v3.4.4) in R (v4.2.1) or seaborn and matplotlib in python.

## Data availability

The single-cell genomics data, including amplicon VCFs, targeted RNA expression and spatial transcriptomics data, will be deposited in GEO. Accession codes will be made available upon publication.

## Code availability

The PRIMR R package for RNA data processing to identify exon-exon junction reads is available from GitHub at https://github.com/Theob0t/PRIMR.

## Acknowledgments

D.A.L. is supported by the Burroughs Wellcome Fund Career Award for Medical Scientists, Valle Scholar Award, Leukemia Lymphoma Scholar Award and the Mark Foundation Emerging Leader Award. This work was also supported by the National Cancer Institute (R33 CA267219), the National Human Genome Research Institute and the Center of Excellence in Genomic Science (RM1HG011014) and the National Institutes of Health Common Fund Somatic Mosaicism Across Human Tissues (UG3NS132139-01), the Japan Agency for Medical Research and Development (AMED): The Core Research for Evolutional Science and Technology (CREST) (JP22gm1110011 to S.O.) and the Moonshot Research and Development Program (JP22zf0127009 to S.O.), AMED (JP23ck0106798 to N.K.), the Japan Society for the Promotion of Science (JSPS), Scientific Research on Innovative Areas (JP15H05909 to S.O.), JSPS KAKENHI (JP20H03660 to A.Y.) and the Japan Science and Technology Agency (JST): Fusion Oriented Research for disruptive Science and Technology (FOREST) Program (JPMJFR215V to N.K.) and Moonshot Research and Development Program (JPMJMS2022-25 to K.N.). This work was made possible by the MacMillan Family Foundation and the MacMillan Center for the Study of the Non-Coding Cancer Genome at the New York Genome Center.

## Author Contributions

D.J.Y., D.A.L., conceived the project, devised the research strategy, and analyzed the data. J.Z., T.B., D.A.L., developed the analytical pipelines for processing the data. A.Y., N.K., S.O. retrieved patient samples for experimental use. D.J.Y., J.Z., T.B., D.A.L. and C.P. wrote the manuscript. D.J.Y., A.H., A.S., J.S., and C.H. performed the experiments. J.Z., T.B., R.R., performed the computational analyses. D.J.Y., J.Z., T.B., D.D., S.W., D.A.L., helped to interpret results. All authors reviewed and approved the final manuscript.

## Supplementary Information

### Supplementary information. Donor metadata

Table 1. Samples used in scRNAseq

Table 2. Samples used in Spatial transcriptomics

Table 3. Samples used in scG2P

### Supplementary Tables

Supplementary table 1. DNA panel genomic locations

Supplementary table 2. RNA panel targets

Supplementary table 3. Cell line mixing variants detected

Supplementary table 4. Comparison with scDNAseq assay

Supplementary table 5. Cell line mixing RNA metadata

Supplementary table 6. Donor samples variants detected

Supplementary table 7. Cell type annotations

Supplementary table 8. RNA module scoring

Supplementary table 9. Donor samples clone call and RNA metadata

## Supplementary Figures

**Supplementary Figure 1.**
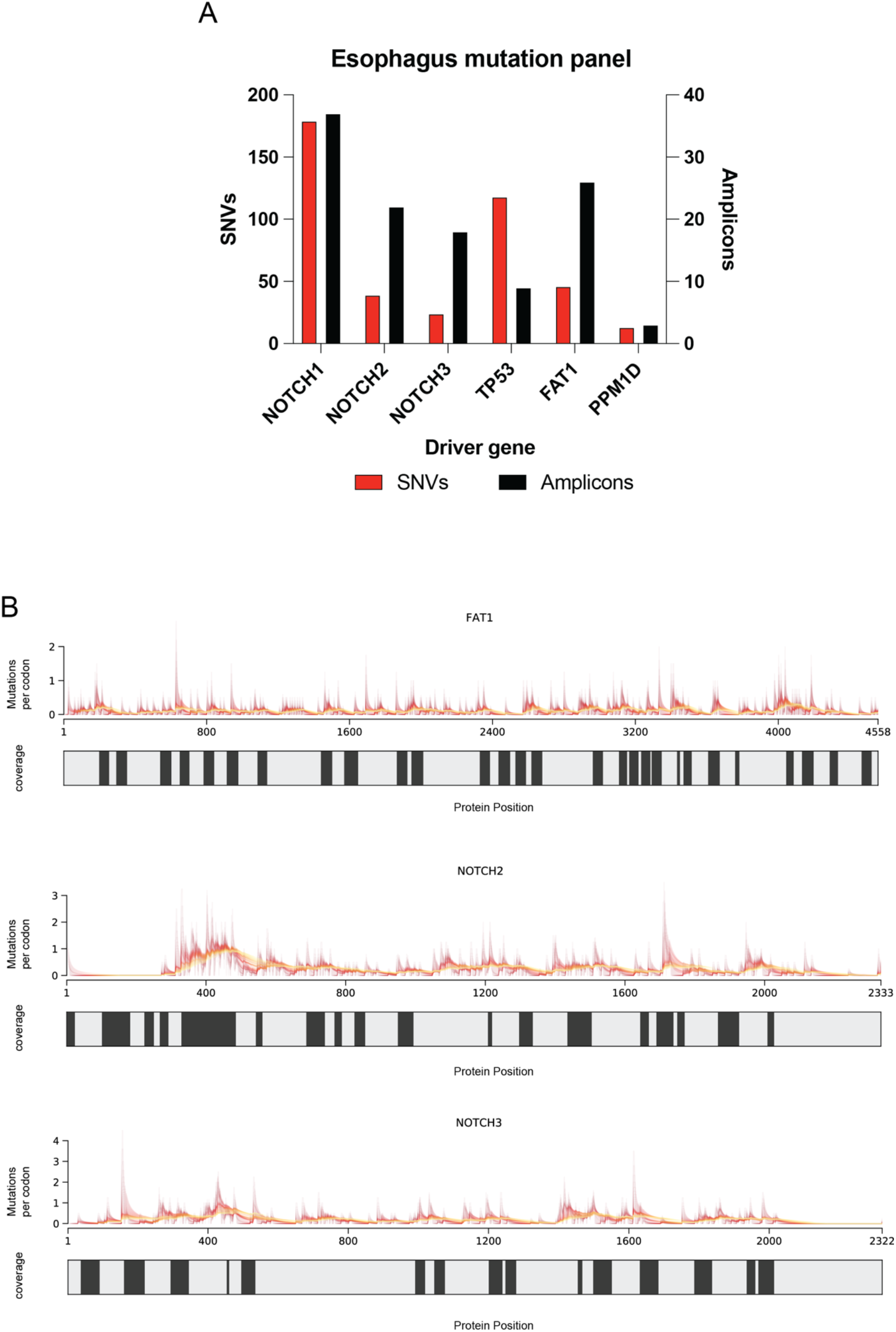
Mutational hot spots in esophageal driver genes. **A**. Summary of the scG2P hotspot mutational panel showing the number of mutations (SNVs) in six driver genes from phenotypically normal esophagus samples (Yokoyama et al.) and number of amplicons designed for each gene. **B**. Amplicon designs for *FAT1, NOTCH2, and NOTCH3*. The number of mutations per codon, as previously reported from bulk sequencing of the esophagus (Yokoyama et al.), is displayed across the protein positions. The color scale represents the size of the rolling window used for calculating mutations per codon, with yellow indicating a 150 bp window and red indicating a 4 bp window. Black bars represent the presence of an amplicon covering the loci.

**Supplementary Figure 2.**
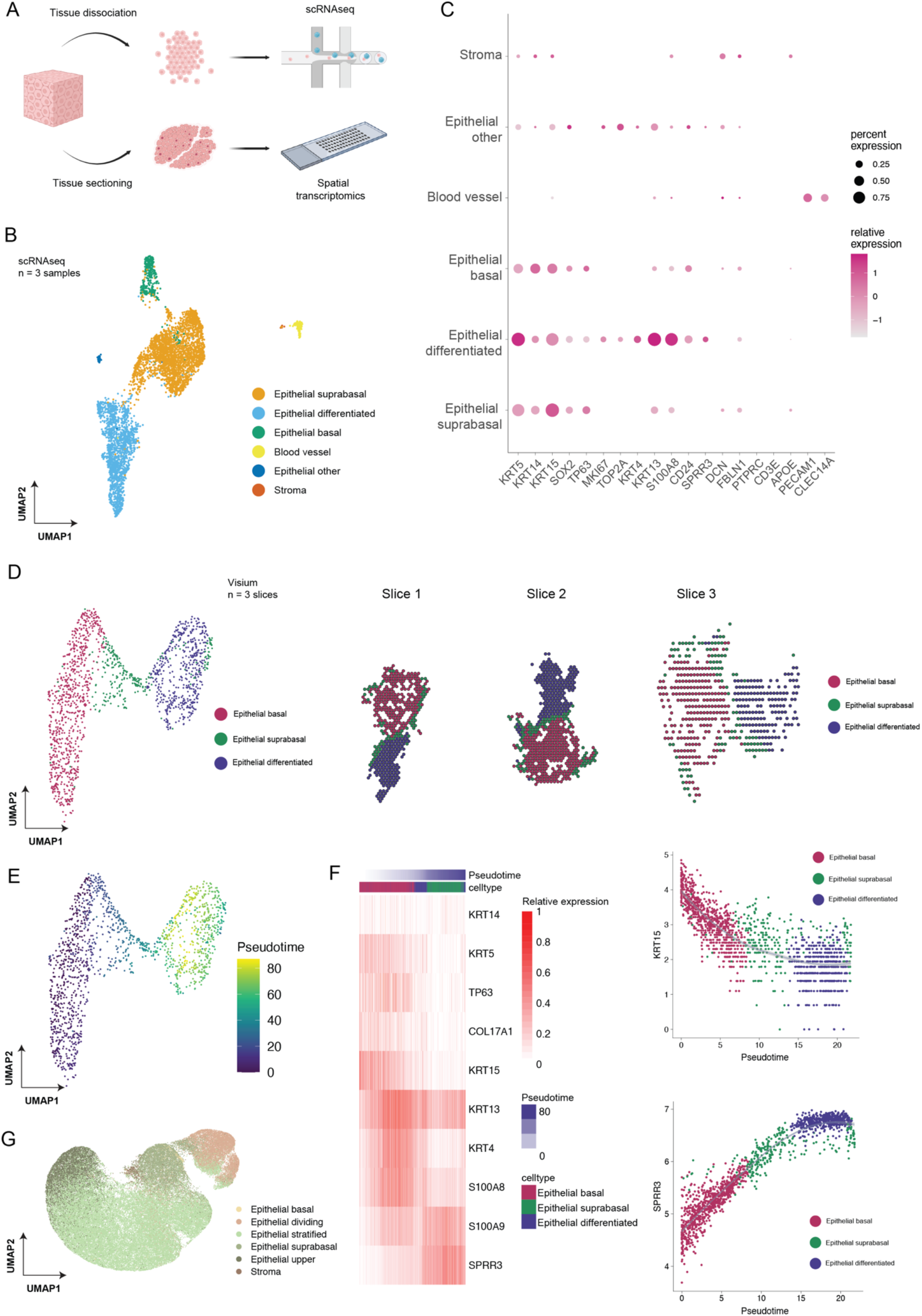
Single-cell RNAseq and spatial transcriptomics of esophageal tissue. **A**. Experimental design for single-cell RNAseq and spatial transcriptomics of esophageal biopsy punches. Punches were dissociated and sorted for EPCAM*+/*CD45-cells for single-cell RNAseq using 10x Chromium assays. Punches were frozen in optimal cutting temperature compound and sectioned for input to the Visium platform for spatial transcriptomics. **B**. UMAP of scRNAseq clustering and cell type annotations from three samples merged (4,912 cells total). Cell type annotations were assigned based on similarity scores to cell type clusters from Madissoon et al. (Methods), capturing stages of differentiation in epithelial cells along with blood vessel and stroma cells. **C**. Expression of gene markers across cell type clusters. Size of dots represent the percentage of cells positive for the marker. Color gradient represents average expression of marker within cell type. **D**. Left, UMAP displaying spatial transcriptomics clustering with cell type annotations based on similarity scores to cell clustering from scRNAseq. Right, spatial projection of cell annotations to microscopy image. Data from 3 slices of sectioned tissue from ESO-ST-3. **E**. Pseudotime of epithelial differentiation from basal epithelial to differentiated epithelial cells. **F**. Left, expression of marker genes with cells organized by pseudotime and annotated by cell type. Right, pseudotime trajectory for expression of basal cell marker gene *KRT15* (top) and differentiated cell marker gene *SPRR3* (bottom). **G**. Cell type clustering and annotations from reference scRNAseq dataset (Madissoon et al.) using only subsetted genes from the scG2P panel. Cell labels are from original author annotations of cell types.

**Supplementary Figure 3.**
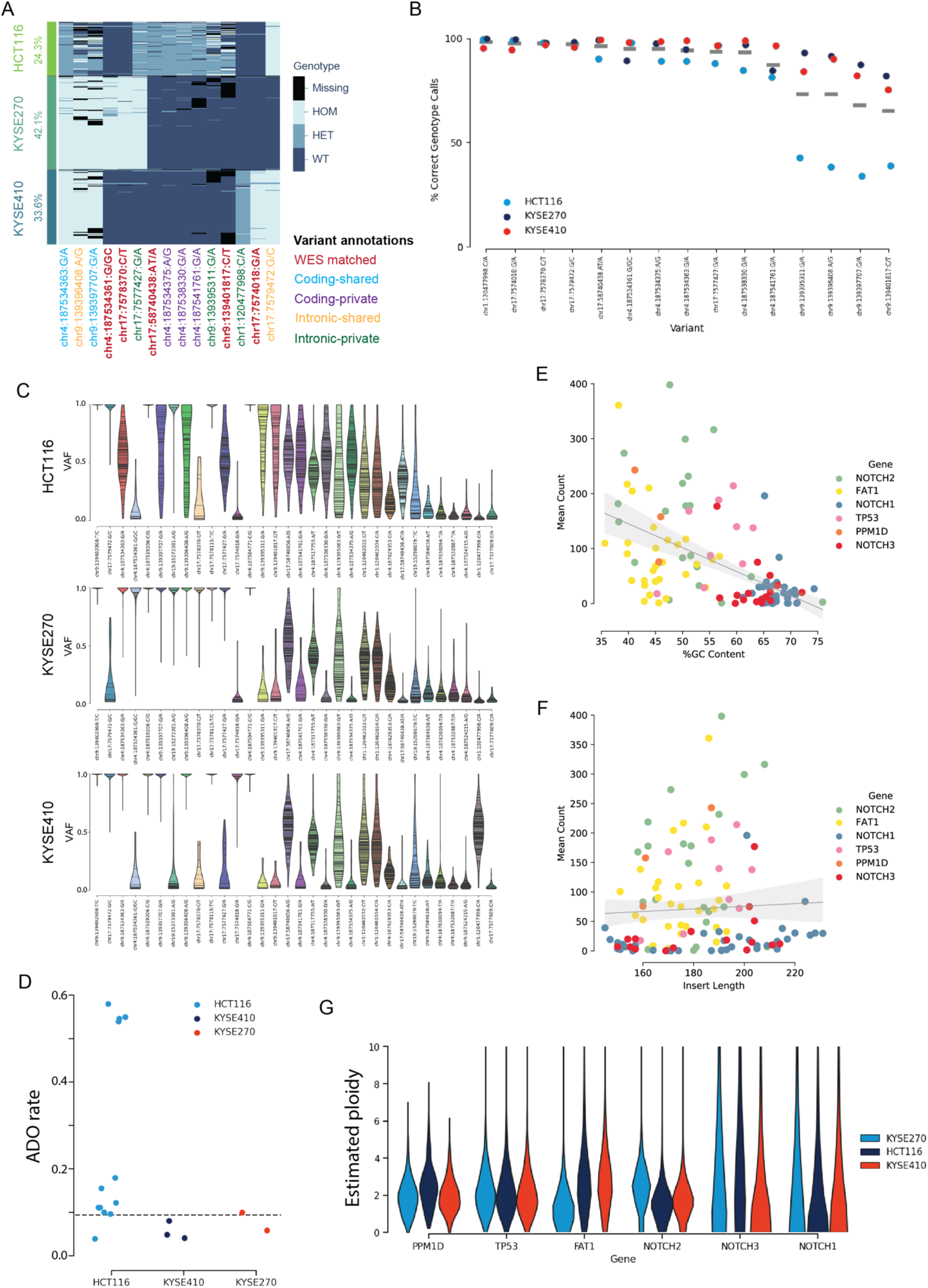
scG2P amplicon performance and variant detection in cell line mixing. **A**. Filtered variants detected in cell line mixing, annotated by type of variant between cell lines. Variant allele frequency reported for variants that are annotated in the GnomAD database. HOM, homozygous; HET, heterozygous; WT, wild-type. **B**. Percentage of cells correctly genotyped for variants detected in whole-exome sequencing (WES) from CCLE in the three cell lines. Line indicates mean accuracy across cell lines. **C**. Distribution of variant allele frequency for filtered variants across cell lines. Each line represents an individual cell. **D**. Allelic dropout (ADO) rate of heterozygous variants detected in cell lines calculated using heterozygous SNPs detected in each cell line. Dashed line represents the median allelic drop out rate across all variants (median = 0.099). **E**. Mean counts of DNA amplicons vs. GC% in cell line mixing experiment, colored by gene. Shaded area represents 95% confidence interval. **F**. Mean counts of DNA amplicons vs. amplicon lengths in cell line mixing experiment, colored by gene. Shaded area represents 95% confidence interval. **G**. Estimated ploidy of driver genes across cell lines calculated using Tapestri copy number detection tool. Depths of coverage of amplicons along a gene was used to infer copy number changes.

**Supplementary Figure 4.**
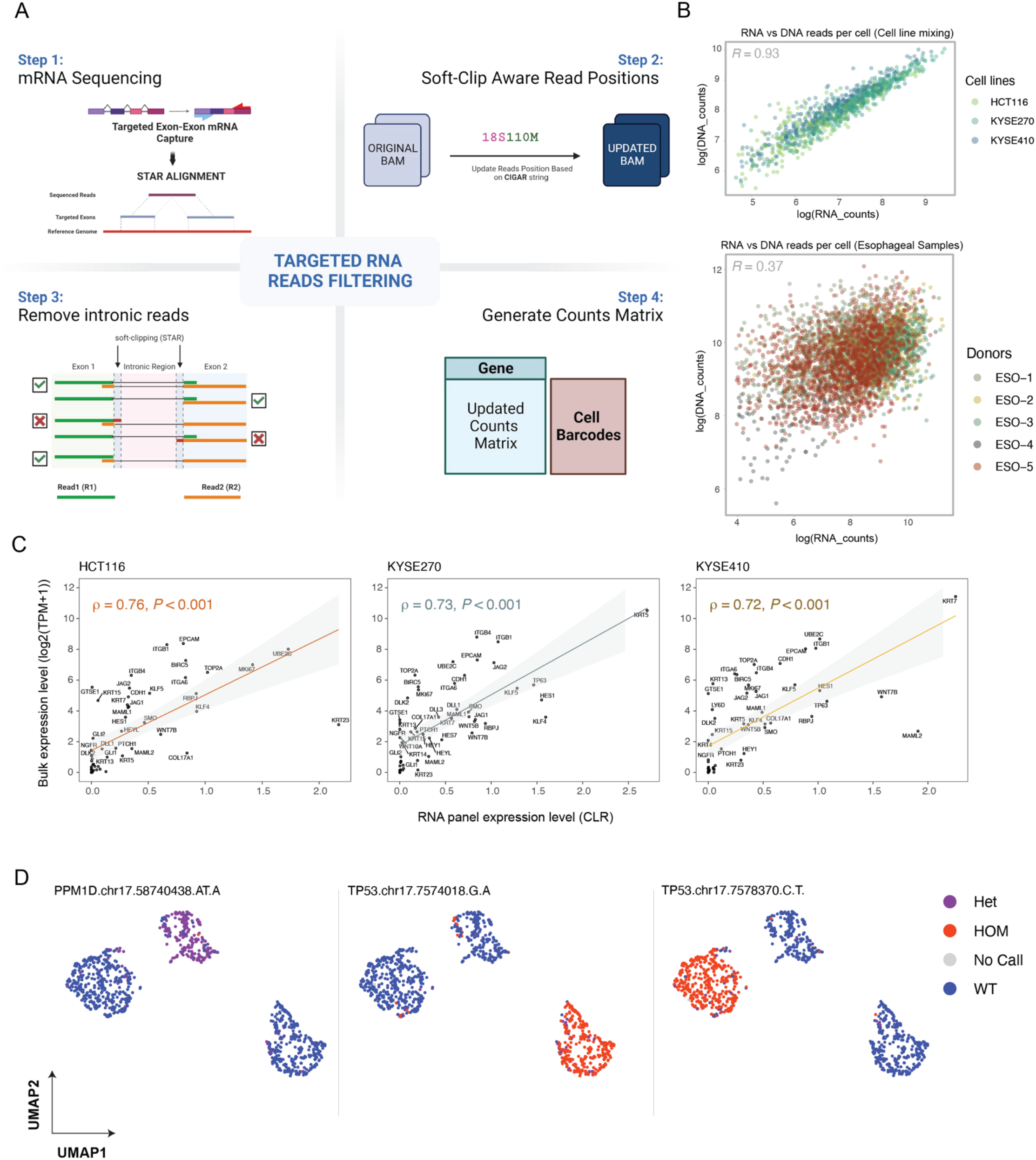
scG2P RNA capture processing and performance. **A**. Schematic of mRNA capture in scG2P. Primers were designed to capture mRNA by targeting exon-exon junctions to distinguish fragments from gDNA and aligned to transcriptome with STAR (Step 1). Aligned BAMs were filtered based on CIGAR scores and soft clipping (Step 2). RNA reads with intronic alignment that had soft clipping on both read 1 and read 2 were filtered (Step 3). Final counts matrix was generated and subsetted based on cell barcodes that passed the DNA processing pipeline (Step 4). **B**. Correlation of captured RNA and DNA reads in cell line mixing (top) and esophageal (bottom) samples. Log of filtered DNA and RNA reads are plotted for cell barcodes called by cell finder pipeline (Methods). **C**. Spearman correlation of bulk RNAseq of cell lines and scG2P RNA panel in cell line mixing. Bulk RNA expression level is measured by log2(TPM+1) from bulk RNAseq downloaded from CCLE. RNA panel expression level is calculated on pseudo-bulk from cell line mixing using centered log ratio (CLR) of counts. Shaded area represents 95% confidence intervals. **D**. Calls for indicated WES variants in each cell line (Fig. 1C) overlaid on RNA-based clustering and UMAP projection. In variants detected from WES data, HCT116 is heterozygous at chr17.58740438.AT.A (left), KYSE410 is homozygous for chr17.5874018.G.A (middle), and KYSE270 is homozygous for chr17.5878370.C.T (right).

**Supplementary Figure 5.**
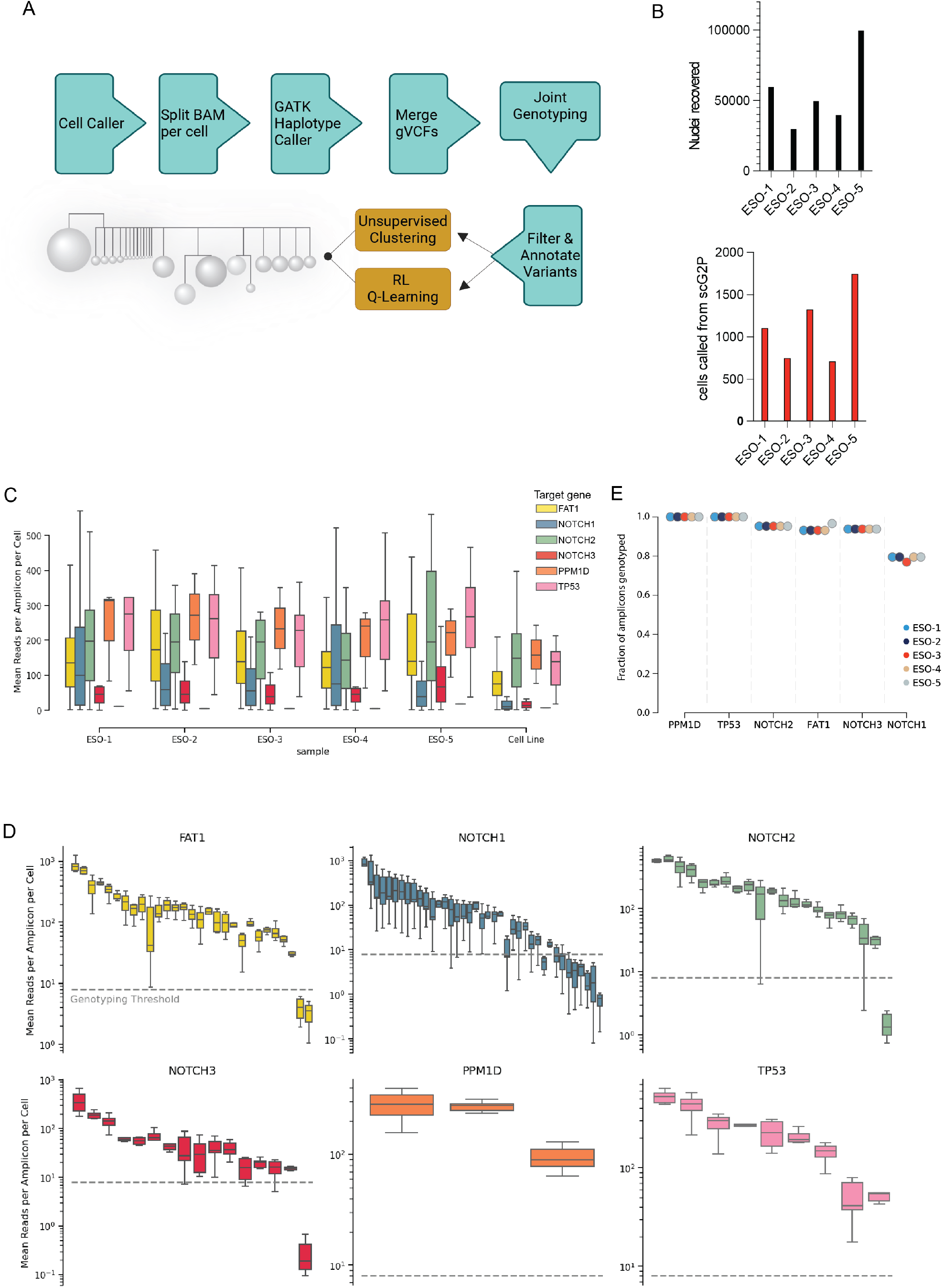
DNA amplicon and genotyping performance. **A**. Workflow for clonal assignment. Cell barcodes are called using coverage metrics utilizing the Tapestri pipeline. Sequences are mapped with BWA Individual aligned BAMs are generated per cell and input to GATK HaplotypeCaller for variant calling. VCFs are generated and merged for join genotyping using VAFs. Variants are filtered based on read depth, genotype quality, and called in 50% of cells. Two clonal inference approaches are used. The first utilizes a soft-clustering of variant allele frequencies. The second utilizes a reinforcement learning model based on Miles et al. **B**. Top, total number of nuclei recovered per pair of esophageal biopsy punches and used as input to assay. Bottom, total number of cell barcodes recovered with cellfinder pipeline. **C**. Amplicon coverage depth, stratified by driver gene, across five esophageal samples and cell line mix (Center line, median; box, IQR; whisker, 1.5*IQR). **D**. Individual amplicons and coverage depth. Dotted line is minimum threshold for genotyping the loci, merged for all esophageal samples (Center line, median; box, IQR; whisker, 1.5*IQR). Amplicons ordered by mean coverage. **E**. Fraction of amplicons in driver genes that had sufficient coverage to be genotyped, across all donor samples. Greater than 80% of all amplicons across all genes were able to be genotyped, with the exception of *NOTCH1* in ESO-3 (75%).

**Supplementary Figure 6.**
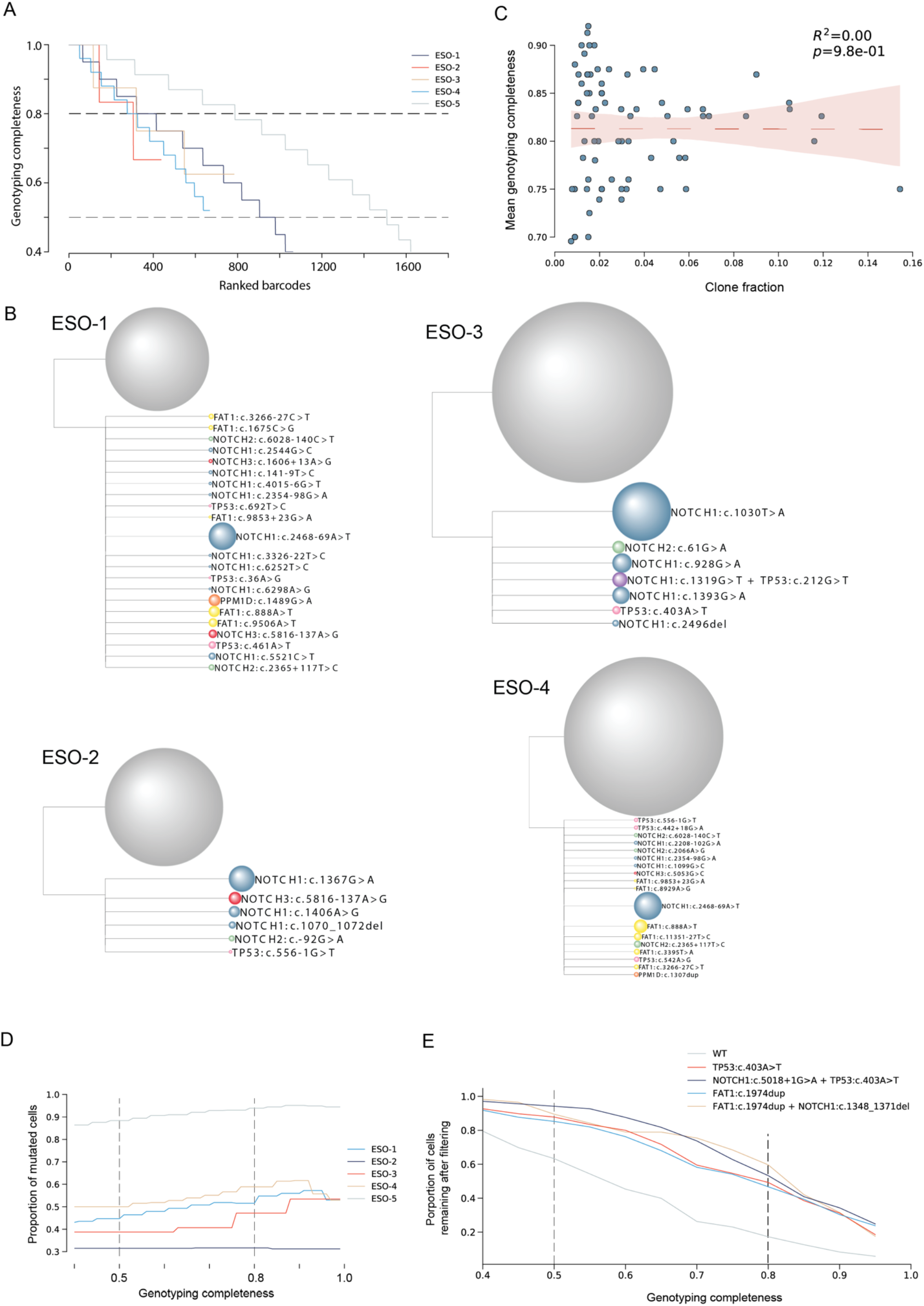
Clonal reconstruction using genotyping completeness. **A**. Number of passing cell barcodes for genotyping completeness filters between 0.40-1.00. Grey dashed lines represent genotyping completeness at 0.50 and 0.80. **B**. Clonal structure of ESO-1,2,3,4. Terminal nodes represent distinct subclones, with node sizes proportional to the relative mutant cell fraction of each subclone. Branch lengths are scaled to reflect the acquisition of a single mutation. **C**. Mean genotyping completeness of cells assigned to a clone vs. clone fraction in all samples. Pearson correlation (R^2^ = 0.00, p-value = 0.98). Dots represent clones detected across all samples. **D**. Fraction of cells called as mutant in each sample when only analyzing cells with genotyping completeness greater than threshold (between 0.40 to 1.00). Dotted lines represent genotyping completeness thresholds of 0.50 and 0.80 as compared in the main text. **E**. Sensitivity analysis of double mutant detection based on genotyping completeness. Proportion of double mutant and parent mutant cells in ESO-5 remaining after filtering at genotyping completeness threshold. Dotted lines represent genotyping completeness thresholds of 0.50 and 0.80 as compared in the main text. Note that the number of cells remaining in double mutant and parent mutant cells are reduced at similar rates as genotyping completeness threshold increases.

**Supplementary Figure 7.**
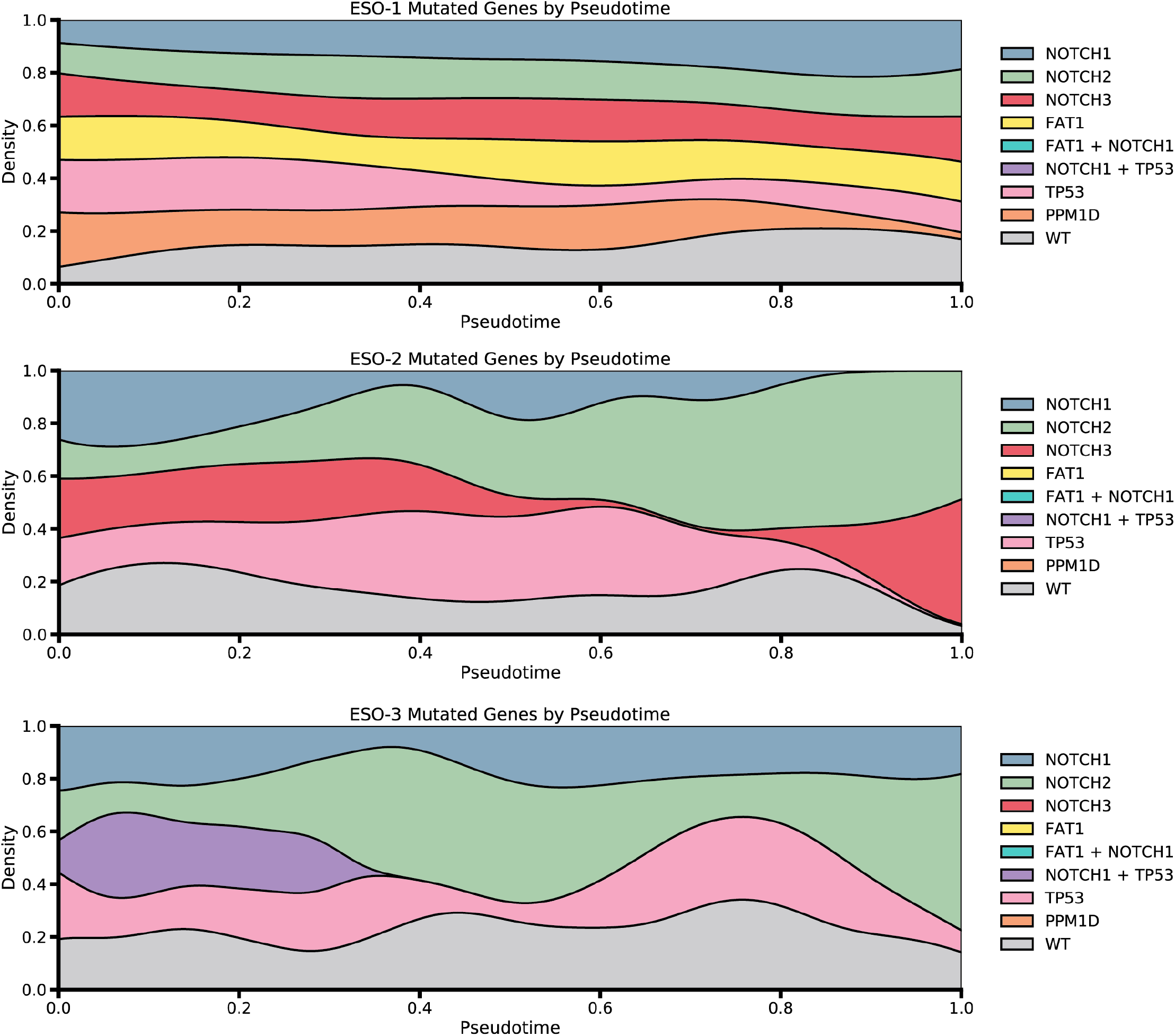
Density of cells with driver mutation across differentiation pseudotime for ESO-1, ESO-2 and ESO-3. Pseudotime from Figure 2C was scaled per donor and binned into deciles to visualize distribution of clones. ESO-4 was not used for diffusion map construction due to the lower number of features detected compared to other samples.

**Supplementary Figure 8.**
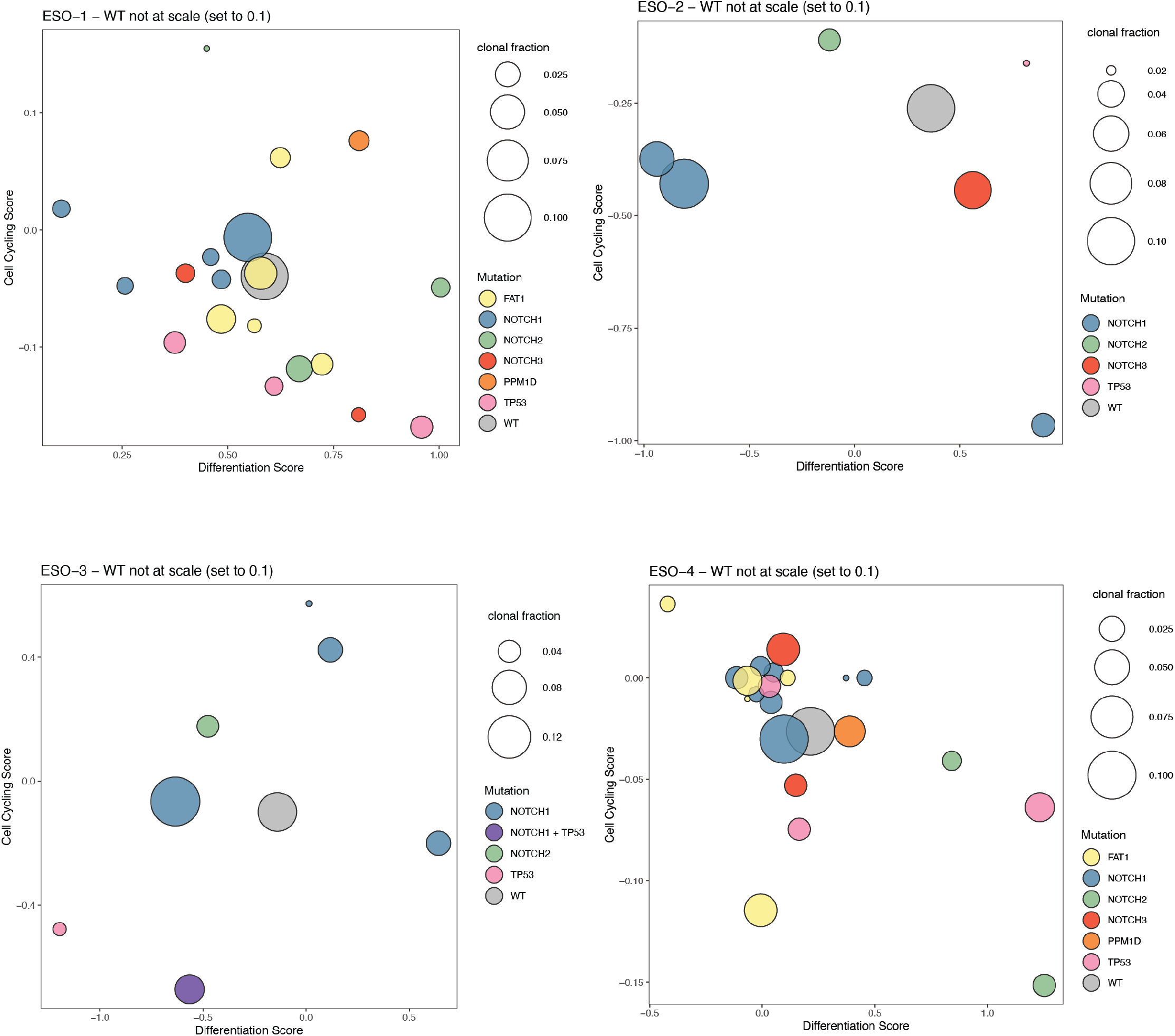
Differentiation and cell cycling scores of clones for ESO-1, ESO-2, ESO-3 and ESO-4. Color indicates mutated gene, size indicates clonal cell fraction. WT cells are fixed at 0.1 fraction size and used to visualize as comparison to clone scores.

**Supplementary Figure 9.**
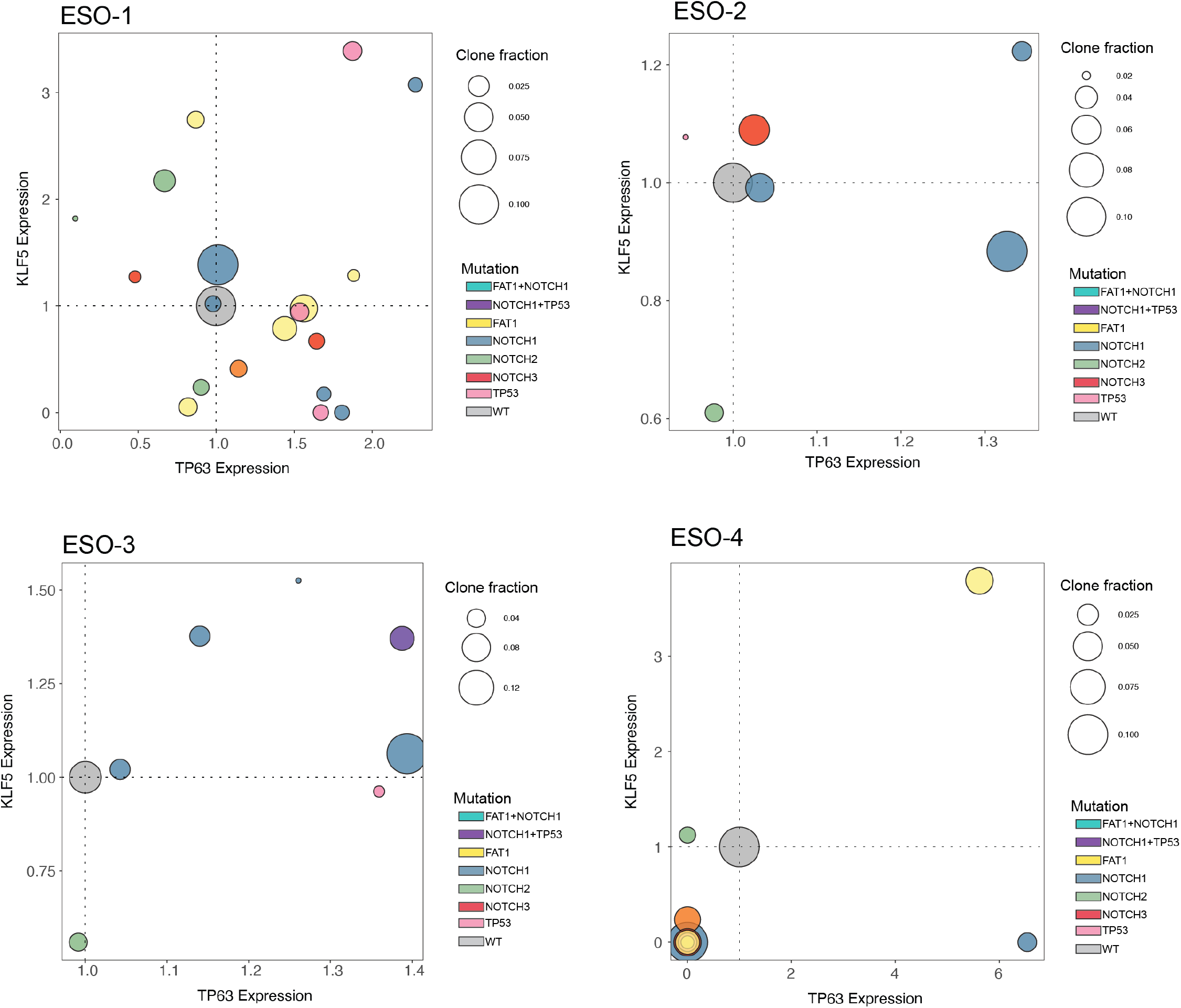
*KLF5* expression versus *TP63* expression of clones from ESO-1, ESO-2, ESO-3 and ESO-4. Color indicates mutated gene, size indicates clonal cell fraction. WT cells are fixed at 0.1 fraction size and used to visualize as comparison to clone scores.

## References

1. Frankell, A. M. et al. The evolution of lung cancer and impact of subclonal selection in TRACERx. Nature 616, 525–533 (2023).

2. Landau, D. A. et al. Evolution and impact of subclonal mutations in chronic lymphocytic leukemia. Cell 152, 714–726 (2013).

3. Landau, D. A. et al. Mutations driving CLL and their evolution in progression and relapse. Nature 526, 525–530 (2015).

4. Martincorena, I. et al. Somatic mutant clones colonize the human esophagus with age. Science 362, 911–917 (2018).

5. Martincorena, I. & Campbell, P. J. Somatic mutation in cancer and normal cells. Science 349, 1483–1489 (2015).

6. Lee-Six, H. et al. Population dynamics of normal human blood inferred from somatic mutations. Nature 561, 473–478 (2018).

7. Nanki, K. et al. Somatic inflammatory gene mutations in human ulcerative colitis epithelium. Nature 577, 254–259 (2020).

8. Olafsson, S. et al. Somatic Evolution in Non-neoplastic IBD-Affected Colon. Cell 182, 672-684.e11 (2020).

9. Anglesio, M. S. et al. Cancer-Associated Mutations in Endometriosis without Cancer. N. Engl. J. Med. 376, 1835–1848 (2017).

10. Ng, S. W. K. et al. Convergent somatic mutations in metabolism genes in chronic liver disease. Nature 598, 473–478 (2021).

11. Brunner, S. F. et al. Somatic mutations and clonal dynamics in healthy and cirrhotic human liver. Nature 574, 538–542 (2019).

12. Yokoyama, A. et al. Age-related remodelling of oesophageal epithelia by mutated cancer drivers. Nature 565, 312–317 (2019).

13. Kakiuchi, N. & Ogawa, S. Clonal expansion in non-cancer tissues. Nat. Rev. Cancer 21, 239–256 (2021).

14. Abby, E. et al. Notch1 mutations drive clonal expansion in normal esophageal epithelium but impair tumor growth. Nat. Genet. 55, 232–245 (2023).

15. Colom, B. et al. Spatial competition shapes the dynamic mutational landscape of normal esophageal epithelium. Nat. Genet. 52, 604–614 (2020).

16. Yin, Y. et al. High-Throughput Single-Cell Sequencing with Linear Amplification. Mol. Cell 76, 676-690.e10 (2019).

17. Macaulay, I. C. et al. G&T-seq: parallel sequencing of single-cell genomes and transcriptomes. Nat. Methods 12, 519–522 (2015).

18. Rodriguez-Meira, A. et al. Unravelling Intratumoral Heterogeneity through High-Sensitivity Single-Cell Mutational Analysis and Parallel RNA Sequencing. Mol. Cell 73, 1292-1305.e8 (2019).

19. Nam, A. S. et al. Somatic mutations and cell identity linked by Genotyping of Transcriptomes. Nature 571, 355–360 (2019).

20. Cortés-López, M. et al. Single-cell multi-omics defines the cell-type-specific impact of splicing aberrations in human hematopoietic clonal outgrowths. Cell Stem Cell 30, 1262-1281.e8 (2023).

21. Miles, L. A. et al. Single-cell mutation analysis of clonal evolution in myeloid malignancies. Nature 587, 477–482 (2020).

22. Morita, K. et al. Clonal evolution of acute myeloid leukemia revealed by high-throughput single-cell genomics. Nat. Commun. 11, 5327 (2020).

23. Nadeu, F. et al. Detection of early seeding of Richter transformation in chronic lymphocytic leukemia. Nat. Med. 28, 1662–1671 (2022).

24. Leighton, J., Hu, M., Sei, E., Meric-Bernstam, F. & Navin, N. E. Reconstructing mutational lineages in breast cancer by multi-patient-targeted single-cell DNA sequencing. Cell Genomics 3, 100215 (2023).

25. Yugawa, T. et al. Regulation of Notch1 Gene Expression by p53 in Epithelial Cells. Mol. Cell. Biol. 27, 3732–3742 (2007).

26. Ohashi, S. et al. NOTCH1 and NOTCH3 coordinate esophageal squamous differentiation through a CSL-dependent transcriptional network. Gastroenterology 139, 2113–2123 (2010).

27. Sakamoto, K. et al. Reduction of NOTCH1 expression pertains to maturation abnormalities of keratinocytes in squamous neoplasms. Lab. Investig. J. Tech. Methods Pathol. 92, 688–702 (2012).

28. Rochman, M. et al. Single-cell RNA-Seq of human esophageal epithelium in homeostasis and allergic inflammation. JCI Insight 7, e159093.

29. Madissoon, E. et al. scRNA-seq assessment of the human lung, spleen, and esophagus tissue stability after cold preservation. Genome Biol. 21, 1 (2019).

30. Busslinger, G. A. et al. Human gastrointestinal epithelia of the esophagus, stomach, and duodenum resolved at single-cell resolution. Cell Rep. 34, (2021).

31. Martincorena, I. et al. High burden and pervasive positive selection of somatic mutations in normal human skin. Science 348, 880–886 (2015).

32. Murai, K. et al. Epidermal Tissue Adapts to Restrain Progenitors Carrying Clonal p53 Mutations. Cell Stem Cell 23, 687-699.e8 (2018).

33. Murai, K. et al. p53 mutation in normal esophagus promotes multiple stages of carcinogenesis but is constrained by clonal competition. Nat. Commun. 13, 6206 (2022).

34. Yang, Y., Goldstein, B. G.Chao, H.-H. & Katz, J. KLF4 and KLF5 regulate proliferation, Apoptosis and invasion in esophageal cancer cells. Cancer Biol. Ther. 4, 1216–1221 (2005).

35. Yang, Y. et al. KLF5 and p53 comprise an incoherent feed-forward loop directing cell-fate decisions following stress. Cell Death Dis. 14, 1–10 (2023).

36. Suliman, Y. et al. p63 Expression Is Associated with p53 Loss in Oral-Esophageal Epithelia of p53-deficient Mice1. Cancer Res. 61, 6467–6473 (2001).

37. Daniely, Y. et al. Critical role of p63 in the development of a normal esophageal and tracheobronchial epithelium. Am. J. Physiol.-Cell Physiol. 287, C171–C181 (2004).

38. Tomasetti, C., Marchionni, L., Nowak, M. A., Parmigiani, G. & Vogelstein, B. Only three driver gene mutations are required for the development of lung and colorectal cancers. Proc. Natl. Acad. Sci. 112, 118–123 (2015).

39. Bailey, M. H. et al. Comprehensive Characterization of Cancer Driver Genes and Mutations. Cell 173, 371-385.e18 (2018).

40. Hu, H. et al. Elevated expression of p63 protein in human esophageal squamous cell carcinomas. Int. J. Cancer 102, 580–583 (2002).

41. Gonzalez-Pena, V. et al. Accurate genomic variant detection in single cells with primary template-directed amplification. Proc. Natl. Acad. Sci. 118, e2024176118 (2021).

42. Nam, A. S., Chaligne, R. & Landau, D. A. Integrating genetic and non-genetic determinants of cancer evolution by single-cell multi-omics. Nat. Rev. Genet. 22, 3–18 (2021).

43. Nam, A. S. et al. Single-cell multi-omics of human clonal hematopoiesis reveals that DNMT3A R882 mutations perturb early progenitor states through selective hypomethylation. Nat. Genet. 54, 1514–1526 (2022).

44. Lee-Six, H. et al. The landscape of somatic mutation in normal colorectal epithelial cells. Nature 574, 532–537 (2019).

45. Ross-Innes, C. S. et al. Whole-genome sequencing provides new insights into the clonal architecture of Barrett’s esophagus and esophageal adenocarcinoma. Nat. Genet. 47, 1038–1046 (2015).

46. Jaiswal, S. et al. Age-related clonal hematopoiesis associated with adverse outcomes. N. Engl. J. Med. 371, 2488–2498 (2014).

47. Yoshizato, T. et al. Somatic Mutations and Clonal Hematopoiesis in Aplastic Anemia. N. Engl. J. Med. 373, 35–47 (2015).

48. Stuart, T. et al. Comprehensive Integration of Single-Cell Data. Cell 177, 1888-1902.e21 (2019).

49. Park, S. R. et al. Single-Cell Transcriptome Analysis of Colon Cancer Cell Response to 5-Fluorouracil-Induced DNA Damage. Cell Rep. 32, 108077 (2020).

50. Wolf, F. A., Angerer, P. & Theis, F. J. SCANPY: large-scale single-cell gene expression data analysis. Genome Biol. 19, 15 (2018).

